# FixNCut: Single-cell genomics through reversible tissue fixation and dissociation

**DOI:** 10.1101/2023.06.16.545221

**Authors:** Laura Jiménez-Gracia, Domenica Marchese, Juan C. Nieto, Ginevra Caratù, Elisa Melón-Ardanaz, Victoria Gudiño, Sara Roth, Kellie Wise, Natalie K Ryan, Kirk B. Jensen, Xavier Hernando-Momblona, Joana P. Bernardes, Florian Tran, Laura Katharina Sievers, Stefan Schreiber, Maarten van den Berge, Tessa Kole, Petra L. van der Velde, Martijn C. Nawijn, Philip Rosenstiel, Eduard Batlle, Lisa M. Butler, Ian A. Parish, Jasmine Plummer, Ivo Gut, Azucena Salas, Holger Heyn, Luciano G. Martelotto

## Abstract

The use of single-cell technologies for clinical applications requires disconnecting sampling from downstream processing steps. Early sample preservation can further increase robustness and reproducibility by avoiding artifacts introduced during specimen handling. We present FixNCut, a methodology for the reversible fixation of tissue followed by dissociation that overcomes current limitations. We applied FixNCut to human and mouse tissues to demonstrate the preservation of RNA integrity, sequencing library complexity, and cellular composition, while diminishing stress-related artifacts. Besides single-cell RNA sequencing, FixNCut is compatible with multiple single-cell and spatial technologies, making it a versatile tool for robust and flexible study designs.

## Introduction

Single-cell sequencing has revolutionized our understanding of the complexity of life, allowing researchers to study tissues, organs and organisms with unprecedented resolution (1). However, most single-cell techniques are designed for freshly prepared specimens, which can present logistical challenges for decentralized study designs that require disconnecting the time and site of sampling from downstream processing steps. In this regard, preservation methods have been developed that enable sample collection and storage, expanding the applications of single-cell sequencing to personalized medicine and collaborative research. In addition to facilitating flexible study designs, early sample preservation can improve robustness and reproducibility by reducing artifacts introduced during sample handling, such as differences in lab personnel skills, library preparation workflows, and sequencing technologies. Furthermore, preservation methods can mitigate cellular stress caused by external factors, such as sample collection, transport, storage, and downstream processing steps involving mechanical or enzymatic dissociation, which can alter transcriptomic profiles (2–4). Such cellular stress can impact sample quality and confound downstream analyses by inducing early stress-response genes and altering the natural state of the cell. Therefore, early sample preservation can enhance the quality and reliability of single-cell sequencing studies, while enabling flexible and decentralized study designs.

Dissociation-induced artifacts can be mitigated by the use of cold active proteases active at low temperatures (6°C) to decrease transcriptional activity and the expression of heat shock and stress-response genes. However, digestion at low temperatures can result in changes in cell type abundance due to incomplete tissue dissociation (2). Alternatively, inhibitors of transcription and translation have been shown to reduce gene expression artifacts by minimizing the impact of dissociation-induced stress (5). To overcome the challenges associated with sample logistics in single-cell studies, cryopreservation has been established as a storage method that preserves transcriptional profiles of cells in suspension and solid tissues (4). However, cryopreservation can result in reduced cell viability and induce considerable changes in sample composition, such as the depletion of epithelial cells, myeloid suppressor cells and neutrophils (3,6–8). Alternatively, cells can be fixed using alcohol, such as ethanol or methanol, but this can cause structural damage due to dehydration, protein denaturation and precipitation, potentially affecting transcriptomic profiles. Nevertheless, alcohol-based fixation has been shown to better maintain cell composition compared to cryopreservation in specific contexts (3,9). More recently, ACME (ACetic-MEthanol) fixation has been developed as a solution to simultaneosly dissociate and fix tissues, resulting in high cellular and RNA integrity (10). Although ACME has been demonstrated to be effective when combined with cryopreservation for cnidarian samples, its value for sample preparation across other species, including mouse and human, remains to be shown. Cross-linking fixatives, such as formaldehyde and paraformaldehyde (PFA), are used with specialized single-cell assays, but are incompatible with commonly applied high-throughput single-cell RNA sequencing (scRNA-seq) protocols. Moreover, formaldehyde-based fixation generally impedes the application of sample multiplexing, single-cell multiome analysis or cellular indexing of transcriptomes and epitopes sequencing (CITE-seq) (10,11). Finally, post-hoc computational tools such as machine learning algorithms have been developed to reduce or remove dissociation-induced artifacts (13). However, due to the often larger biological compared to technical variability and the fact that not all cell types within a sample suffer the same stress, it is difficult to generalize bias correction across cells and samples (14).

To overcome the challenges discussed above, it was crucial to develop a standardized protocol for sample collection, cell stabilization, and tissue processing that allows for fixation prior to sample processing. We present a workflow, called FixNCut, which uses Lomant’s Reagent (dithiobis(succinimidyl propionate), DSP), a reverse crosslinker fixative, to enable tissue fixation prior to sample digestion steps. Such order of events prevents changes in gene expression during digestion and processing, while disconnecting sampling time and location from subsequent processing sites. DSP is a homo-bifunctional N-hydroxysuccinimide ester (NHS ester) crosslinker that contains an amine-reactive NHS ester at each end of an 8-carbon spacer arm. NHS esters react with primary amines at pH 7-9 to form stable amide bonds and release N-hydroxy-succinimide. Proteins typically have multiple primary amines in the side chain of lysine residues and the N-terminus of each polypeptide that can serve as targets for NHS ester crosslinking reagents. DSP is lipophilic and membrane-permeable, making it applicable for intracellular and intramembrane conjugation. However, it is insoluble in water and must be dissolved in an organic solvent before adding to the reaction mixture. The presence of a disulfide bond in the center of DSP makes its crosslinking reversible via reducing agents like DTT, which is present in most reverse transcription buffers of single-cell sequencing applications (e.g. 10x Genomics assays). To date, DSP has been successfully used to preserve cells for single-cell sequencing in applications such as CLint-Seq, nanofluidic systems or phosphoprotein-based cell selection (15–17).

Here, we demonstrate the versatility of the FixNCut protocol to overcome key limitations in generating single-cell data. We provide evidence that FixNCut preserves RNA integrity, library complexity, and cellular composition, while allowing for cell labeling or sample hashing prior to single-cell analysis. To illustrate its potential, we applied FixNCut to fix and digest mouse lung and colon tissue, as well as human colon biopsies from inflammatory bowel disease (IBD) patients, demonstrating its clinical utility. Additionally, we show that DSP fixation can be used in the context of spatial-omics, specifically multiplexing tissue imaging for spatial proteomics (i.e. Phenocycler).

## Results

### Reversible fixation of human cells in suspension

Extending previous studies using DSP to preserve cell lines for RNA sequencing (15), we initially confirmed its applicability for single-cell analysis of cells in suspension (human peripheral blood mononuclear cells; PBMCs) and microfluidics systems (10x Genomics Chromium Controller), before combining fixation and dissociation of complex solid tissues. To this end, we compared cell morphology, RNA integrity, and reverse transcription efficiency of fresh and DSP-fixed PBMCs. Fixed cells showed highly similar morphology compared to fresh PBMCs in bright-field microscopy, with no evident changes in cell phenotypes or sample clumping after fixation (Fig. S1a). Next, we captured and barcoded single cells from both fresh and DSP-fixed samples using the Next GEM Single Cell 3’ Reagent Kits v3.1 from 10x Genomics. Bioanalyzer profiles of the amplified cDNA from both samples were virtually identical, demonstrating DSP fixation not to affect RNA integrity or reverse transcription performance (Fig. S1b,c). After sequencing, we confidently mapped over 80% of the reads from both sequencing libraries to the human reference genome, with over 50% of exonic reads usable for quantifying gene expression levels (Fig. 1a). We observed a comparable correlation between the number of detected genes and sequencing depth for fresh and fixed samples, indicating DSP fixation to conserve library complexity (Fig. 1b). We confirmed this observation at the single-cell level, where we found a similar relationship between sequencing depth and the number of detected unique molecular identifiers (UMIs) or genes per cell (Fig. S1d). In line, we observed similar gene counts in single blood cells in fresh and fixed samples (Fig. 1c). After filtering out low quality cells, we found a similar distribution of the main quality control (QC) metrics between both protocols (Fig. S1e), except for a few specific cell subpopulations (Fig. S1f). These results suggest DSP fixation to conserve the ability to detect gene transcripts in single cells compared to fresh samples in scRNA-seq experiments.

**Figure 1.**
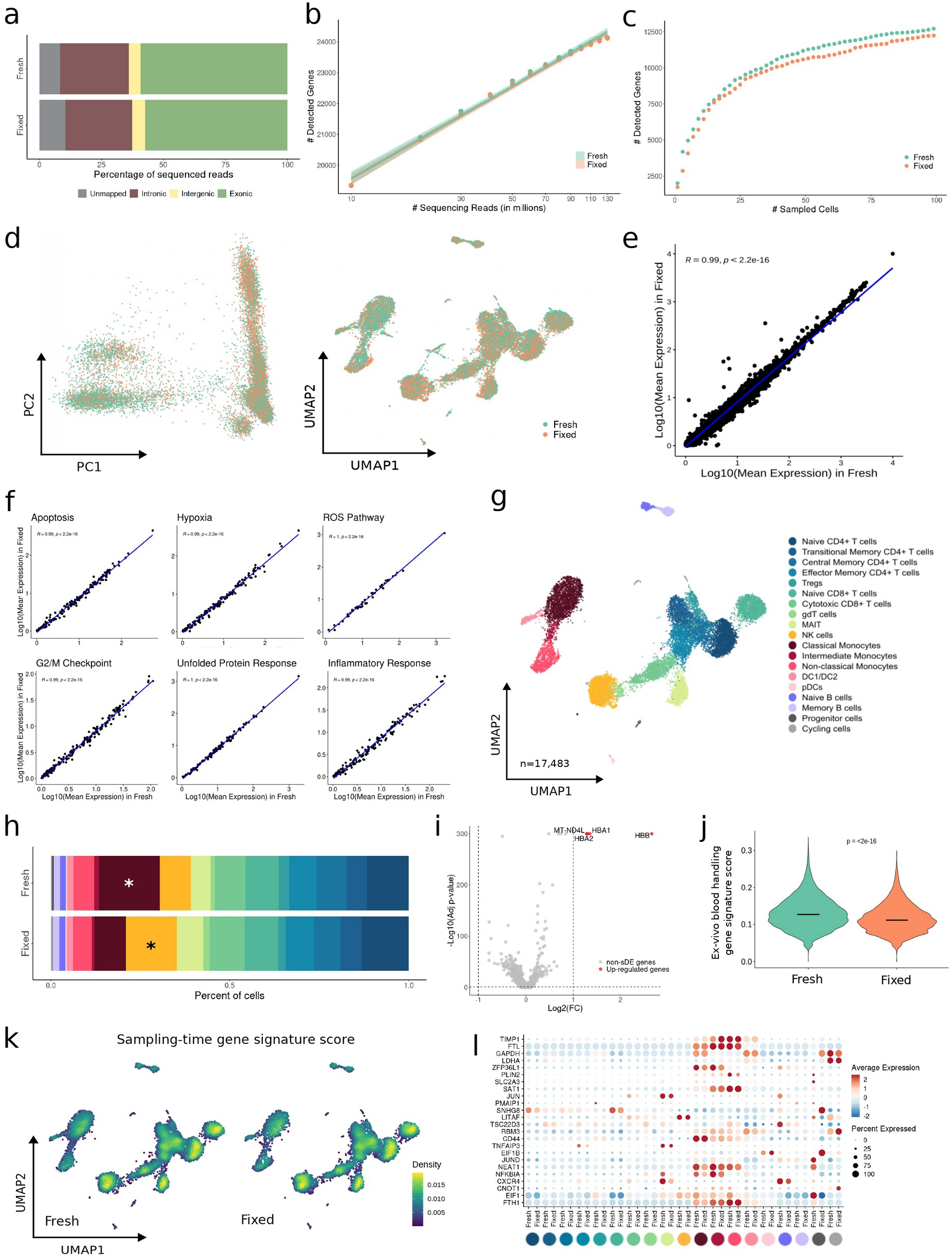
FixNCut protocol in human peripheral blood mononuclear cells (PBMCs). (**a**) Mapping analysis of sequencing reads within a genomic region. (**b**) Comparative analysis of the number of detected genes based on sequencing reads using a linear model. (**c**) Cumulative gene counts analyzed using randomly sampled cells. (**d**) Principal component analysis (PCA) and Uniform Manifold Approximation and Projection (UMAP) representation of gene expression profiles variances of fresh and fixed samples. (**e-f**) Linear regression model comparing average gene expression levels of expressed genes (**e**) and main biological hallmarks, including apoptosis, hypoxia, reactive oxygen species (ROS), cell cycle G2/M checkpoint, unfolded protein response (UPR), and inflammatory response genes (**f**). The coefficient of determination (R2) computed with Pearson correlation and the corresponding p-value are indicated. (**g**) UMAP visualization of 17,483 fresh and fixed PBMCs, colored by 19 cell populations. (**h**) Comparison of cell population proportions between fresh (n=9,754) and fixed (n=7,729) PBMCs with the Bayesian model scCODA. Asterisks (*) indicate credible changes. (**i**) Differential gene expression analysis between fresh and fixed samples. The top differentially expressed genes (DEGs) with significant adjusted p-values (FDR) < 0.05, up-regulated (red), and down-regulated (blue) with Log2FC > 1 are indicated. (**j**) Violin plot of ex-vivo blood handling gene signature score (18) for fresh and fixed human PBMCs. Statistical analysis between fixed and fresh cells was performed using the Wilcoxon signed-rank test. (**k**) UMAP visualization of sampling-time gene signature score (4) for fresh and fixed human PBMCs. (**l**) Dotplot showing the average expression of sampling-time DEGs for Fresh (y-axis) for all 19 cell types (x-axis) split by protocol. The dot size reflects the percentage of cells in a cluster expressing each gene, and the color represents the average expression level.

To further assess potential technical variation between protocols, we identified highly variable genes (HVGs) independently in fixed and fresh PBMCs. We found that 70% of HVGs were shared between the two protocols, indicating a conserved representation of the transcriptome and suitability for joint downstream processing (Fig. S1g). Additionally, when we examined the variation captured in the main principal components (PCs) and displayed single-cell transcriptomes in two dimensions (uniform manifold approximation and projection; UMAP), we did not observe any notable outliers due to the sampling protocol (Fig. 1d). Cells clustered together based on biological differences rather than preparation protocol, suggesting fixed and fresh cells to have similar capacity for cellular phenotyping. The pseudo-bulk gene expression profiles between fixed and fresh samples were highly correlated (R2 = 0.99, p < 2.2e-16) (Fig. 1e), indicating DSP fixation not to alter the expression of specific genes. This was further confirmed at the cell population level (Fig. S1h). Moreover, the biological processes such as apoptosis, hypoxia, reactive oxygen species (ROS), cell-cycle (G2/M checkpoint), unfolded protein response (UPR), and inflammation hallmarks remained unchanged across libraries (Fig. 1f).

Next, we performed joint analysis of 17,483 fresh and fixed cells, which were clustered to define 19 distinct cell populations (Fig. 1g; Fig. S1i). All cell types and states were found across both protocols at similar proportions, except for classical monocytes and NK cells, which showed small but significant differences, being slightly increased in fresh and fixed, respectively (Fig. 1h). Fixation did not affect differential expression analysis (DEA), with only four up-regulated genes representing hemoglobin subunits (HBA1, HBA2, and HBB) and a mitochondrial gene (MT-NDL4) (Fig. 1i; Additional file 2: Table 1). These genes were consistently found across all cell populations (Additional file 3: Table 2), a phenomenon also observed when performing digestion protocols at low temperatures (3). The FixNCut protocol may prevent erythrocyte lysis, leading to their co-encapsulation with nucleated blood cells and detection of specific transcripts.

**Table 2:** Recommendations on working with DSP stock and working dilution, as described by Attar *et al.* (15)

|  |
| --- |
| **Preparation of 50X DSP stock** |
| ● Equilibrate DSP vial at RT for 30 min and then prepare a 50X stock solution of DSP (50 mg/mL) in anhydrous dimethyl sulfoxide (Sigma, Cat. N. 276855-100ML).<br>● Dispense the stock into single use aliquots (e.g. 100 µL aliquots, but volume depends on your use) and store in a bag & dry environment (with silica/desiccant if possible) at -80°C.<br>● Be mindful of not opening and closing the tubes at -80°C. |
| **Preparation of DSP working dilution** |
| ● Thaw the 50X DSP stock reagent from the -80°C and equilibrate at RT no longer than 10 min before fixation.<br>● Immediately before use prepare 500 µL of 1X DSP working solution in molecular biology grade 1X PBS as follows: aliquot 10 µL of 50X DSP stock reagent in a 1.5 mL tube and while vortexing (VERY IMPORTANT) add 490 µL of PBS dropwise using a P200 pipette. 1X DSP must be used within 5 min of preparation.<br>● *Note: Do not prepare larger volumes (e.g. if you need to fix two samples, prepare each aliquot of 500 µL separately, DO NOT prepare 1 mL and then aliquot into two tubes). You should notice some thin white rings on the walls of the tube once diluted the 50X stock. This is expected and will be cleared during filtration. Stronger precipitation indicates insufficient solving of DSP and preparation of a new dilution is strongly recommended.*<br>● Filter the 1X DSP working dilution using a 40 µm Flowmi strainer (Sigma, cat. no. BAH136800040-50EA). After filtration 1X DSP can be pooled.<br>● Do not re-freeze the leftovers of the 50X DSP. Always use a freshly thawed aliquot to prepare the 1X working solution. |

Importantly, we observed a reduction in technical artifacts introduced during sample processing prior to single-cell experiments upon fixation. Specifically, gene expression alterations previously defined to correlate with *ex vivo* PBMC handling (18) showed a significant reduction upon fixation (p < 2.2e-16) (Fig. 1k). Moreover, a sampling-time gene signature obtained from single-cell benchmarking studies (4) also showed a significant reduction in the fixed PBMCs (p < 2.2e-16) (Fig. 1j). Interestingly, T lymphocytes appeared to be particularly affected (p <= 0.0001), showing the strongest protection from sampling artifacts in fixed cells (Fig. S1j). Main driver genes of such variation included EIF1, CXCR4, NFKBIA, NEAT1, JUND, SNHG8, and JUN (Fig. 1l). DSP also protected against the general reduction of gene expression activity, previously reported during PBMC sample processing (4). Notably, more than 30% of genes from the sampling-time signature were also detected as enriched in the fresh PBMCs (Fig. S1k). The results suggest that fixed PBMCs have comparable cellular composition and gene expression profiles to freshly prepared samples, while reducing gene expression artifacts introduced during sample preparation.

### FixNCut protocol applied on mouse solid tissues

Beyond the benefits of cell fixation in standardizing sample processing and preserving gene expression profiles of cells in suspension, the FixNCut protocol was specifically designed for solid tissues. Specifically, it allows for fixation and subsequent digestion, which is particularly advantageous for complex and logistically challenging study designs, such as clinical trials. Here, sampling artifacts, including biases in gene expression and cell type composition, are frequently observed in fragile solid tissue types. Fixation prior to digestion using the FixNCut protocol can critically reduce these artifacts. Thus, we next evaluated the effectiveness of the FixNCut protocol with subsequent scRNA-seq readout in different solid mouse tissues before extending its application to challenging human patient samples, such as tissue biopsies.

Fresh mouse lung samples were minced, mixed, and split into two aliquots, one processed fresh and the other fixed using the FixNCut protocol with subsequent 30 minute tissue digestion using Liberase TL. Experiments were conducted in biological replicates. The fixed sample showed a slight decrease in cell size and an increase in DAPI+ cells, but overall, cell morphology was similar to the fresh sample (Fig. S2a). Single-cell encapsulation and scRNA-seq (10x Genomics, 3’ RNA v3.1) showed comparable proportions of reads mapped to the mouse reference genome and exonic genomic regions for both fresh and fixed samples (Fig. 2a). We further observed a similar correlation between the number of detected genes and sequencing depth in fresh and fixed samples (Fig. 2b). Notably, at the same sequencing depth, fixed lung samples showed more detected genes in both replicates, mostly lncRNAs, indicating higher library complexities (Fig. S2b). Our analysis of the scRNA-seq data from fresh and fixed lung samples revealed that genes detected in all samples displayed higher expression values for fixed (median of Mean Expression (ME) = 0.0314) as compared to genes detected in fresh (median ME = 0.0244) (Additional file 4: Table 3). This was also observed for the fixed-exclusive captured genes (median ME = 0.1642) compared to fresh-exclusive (median ME = 0.1647), indicating that FixNCut enhances the gene capture efficiency, potentially allowing for a more fine-grained cell phenotyping after sample fixation. At the cell level, we confirmed the higher complexity of fixed libraries, as reflected by the number of detected UMIs and genes (Fig. S2c). We further observed that more genes were captured for the fixed samples after accumulating information from individual lung cells (Fig. 2c). Importantly, after filtering out low-quality cells and neutrophils, the main QC metrics in fixed samples showed more consistent distributions across all characterized cell types (Fig. S2d,e), suggesting that FixNCut protocol not only preserves the capacity for scRNA-seq profiling after fixation and digestion, but further improves the detection of gene transcripts.

**Figure 2.**
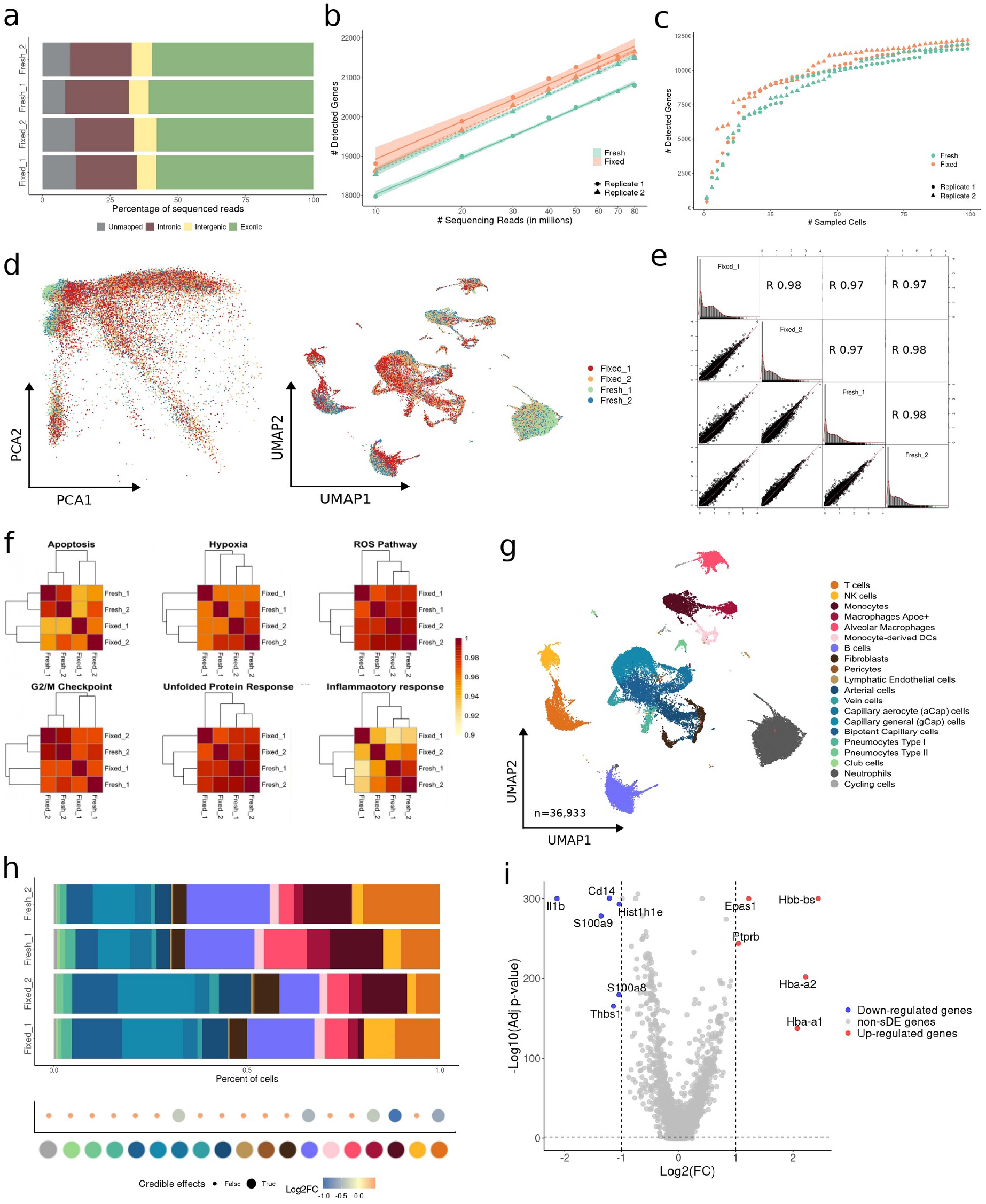
FixNCut protocol tested in mouse lung samples. (**a**) Mapping analysis of sequencing reads within a genomic region. (**b**) Comparative analysis of the number of detected genes based on sequencing reads using a linear model. (**c**) Cumulative gene counts analyzed using randomly sampled cells. (**d**) Principal component analysis (PCA) and Uniform Manifold Approximation and Projection (UMAP) representation of gene expression profile variances of fresh and fixed replicated samples. (**e**) Linear regression model comparing average gene expression levels of expressed genes across protocol and replicates. The coefficient of determination (R2) computed with Pearson correlation is indicated. (**f**) Hierarchical clustering of coefficient of determination (R2) obtained for all pair comparisons across protocol and replicates for biological hallmarks, including apoptosis, hypoxia, reactive oxygen species (ROS), cell cycle G2/M checkpoint, unfolded protein response (UPR), and inflammatory response genes. (**g**) UMAP visualization of 36,933 fresh and fixed mouse lung cells, colored by 20 cell populations. (**h**) Comparison of cell population proportions between fresh (n1 = 3813 and n2 = 6696) and fixed cells (n1 = 10704 and n2 = 6974). The top figure shows the difference in cell population proportions between fresh and fixed replicated samples, and the bottom figure shows the results of the compositional cell analysis using the Bayesian model scCODA. Credible changes and Log2FC are indicated. (**i**) Differential gene expression analysis between fresh and fixed samples. The top differentially expressed genes (DEGs) with significant adjusted p-values (FDR) < 0.05, up-regulated (red), and down-regulated (blue) with Log2FC > 1 are indicated.

**Table 3:** Flow Cytometry Antibodies and Reagents.

| Antibody | Clone | Conjugate | Company |
| --- | --- | --- | --- |
| Human BD Fc Block | Fc1 | NA | BD # 564219 |
| Anti-human CD45 | HI30 | BV510 | BD # 563204 |
|  | HI30 | PE | BD Pharmingen |
| Anti-human CD3 | UCHT1 | BV711 | BD # 563725 |
|  | SK7 | PE-Cy7 | Biolegend PN-344816 |
|  | OKT3 | PerCP | Biolegend |
| Anti-human CD4 | SK3 | BV650 | BD # 563875 |
|  | RPA-T4 | Alexa Fluor® 488 | Biolegend PN-300519 |
| Anti-human CD8 | HIT8a | PE | BD #555635 |
|  | SK1 | APC-Cy7 | Biolegend PN-344714 |
| Anti-human CD19 | 6E10 | Alexa Fluor® 647 | Biolegend PN-302222 |
| Anti-human CD298 | LNH-94 | PE-Cy7 | BioLegend #341707 |
| Anti-human β2-microglobulin | 2M2 | APC | BioLegend #316311 |
| anti-human CD11b | ICRF44 | PeCy7 | Biolegend |
| anti-human EPCAM | 9C4 | APC | Biolegend |
| Annexin V |  | FITC | BD #556420 |
| Propidium Iodine |  |  | Sigma-Aldrich #P4864 |

An overlap of almost 60% of sample-specific HVGs was found across all four samples, which increased to 80% when directly comparing the fresh and FixNCut protocols (Fig. S2f). The absence of batch-effects linked to replicates or protocols was demonstrated by the PCA and UMAP representations (Fig. 2d; Fig. S2g), indicating bias-free transcriptome profiles after cell fixation and digestion. Highly comparable profiles of mean gene expression values were observed between fresh and fixed mouse lung samples (all comparisons, R2 > 0.97, p < 2.2e-16) (Fig. 2e), a finding also confirmed at the population level (Fig. S2h). Moreover, the high correlation across gene programs supported the absence of alterations in major biological processes (Fig. 2f).

We then performed a joint analysis of all 36,933 mouse lung cells, which segregated into 20 distinct cell populations, encompassing both lung and tissue-resident immune cells (Fig. 2g; Fig. S2i). All characterized cell types were detected in both fresh and fixed samples. However, we observed a notable variation in the neutrophil fraction between replicates. To account for the differences in mouse collection protocol between replicate samples, we excluded neutrophils from downstream analysis. When analyzing the remaining 28,187 cells, we observed variability in cell type proportions between both protocols. The fixed protocol showed an improved representation of tightly connected epithelial and endothelial cell types, while immune cells (B and T cells, monocytes, and macrophages Apoe+) and aCap cells were proportionally reduced (Fig. 2h). To validate preserved gene expression profiles in fixed tissues, we performed differential expression analysis (DEA) between the two protocols. We observed up-regulation of genes related to hemoglobin subunits (Hbb-bs, Hba-a2, and Hba-a1), hypoxia (Epas1), and endothelial cell markers (Ptprb) for the fixed protocol, which could be largely explained by the enrichment of this population upon fixation. In contrast, fresh samples were enriched in genes related to inflammatory and immune processes and Hist1h1e, in accordance with the increased proportion of recovered immune cells (Fig. 2i; Additional file 5: Table 4). Consistent with observations for PBMCs, we found a uniform enrichment in the hemoglobin genes for the fixed sample across cell types (Additional file 6: Table 5). Gene set enrichment analysis (GSEA) stratified by cell type revealed that freshly prepared endothelial cells were enriched in ROS, apoptosis and cellular response to external stimuli, whereas the opposite patterns were observed upon fixation (Additional file 7: Table 6). Overall, these results suggest the global conservation of library complexity and quality, along with improved metrics for gene detection and the inclusion of tightly connected, challenging-to-isolate cell types in fixed mouse lung samples.

**Table 4:**
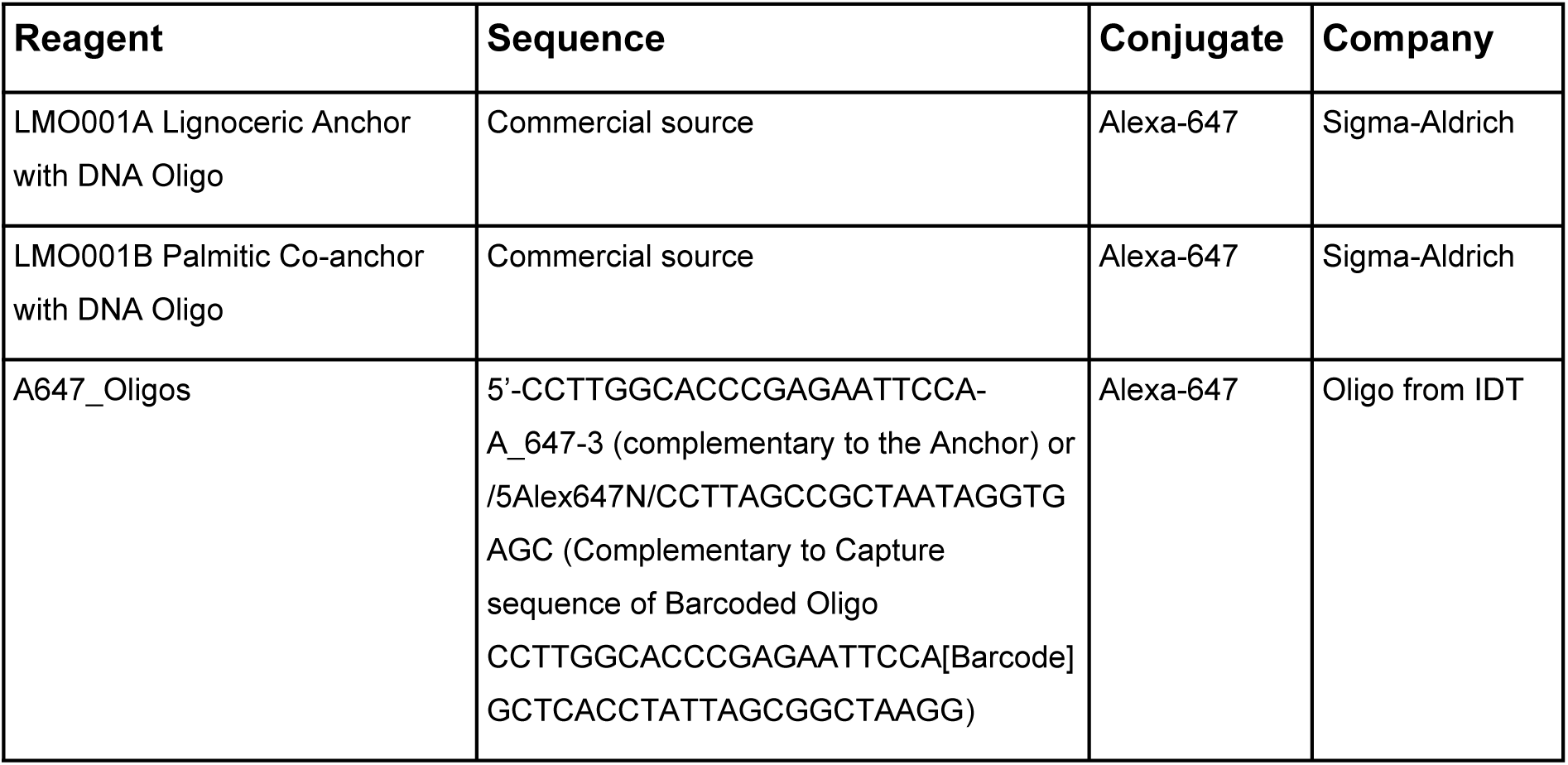
Flow Cytometry Lipid Modified Oligos (LMOs)

**Table 5:** Antibody and reporter cycle layout, including LED intensities and exposure times for each marker.

| Cycle # | Cy3<br>(LED 100%) | Exposure time (ms) | Cy5<br>(LED 100%) | Exposure time (ms) | Cy7<br>(LED 60%) | Exposure time (ms) |
| --- | --- | --- | --- | --- | --- | --- |
| 1 | BLANK | 250 | BLANK | 350 | BLANK | 500 |
| 2 | e-cadherin-RX014 | 750 | ERG-RX031 | 850 | SMA-RX013 | 500 |
| 3 | CD8-RX026 | 750 | CD3e-RX045 | 850 | PanCK-RX019 | 400 |
| 4 | Empty | 250 | p63-RX046 | 700 | Empty | 500 |
| 5 | BLANK | 250 | BLANK | 350 | BLANK | 500 |

We further evaluated the performance of the FixNCut protocol in a different challenging solid tissue context. To do so, we minced and mixed mouse colon samples that were split and subjected to scRNA-seq after digestion of either fresh or fixed tissues. Our results indicate that FixNCut provides several benefits, including improved transcriptome capture accuracy, as evidenced by a higher number of total reads mapped to the reference and a higher exonic fraction (Fig. 3a). Additionally, the fixed sample exhibited a considerably higher library complexity (Fig. 3b), with fixed cells showing increased numbers of detected UMIs and genes (Fig. S3a). Notably, the cumulative gene count was greater for the fixed colon (Fig. 3c), and we observed improved QC metrics for the FixNCut sample after filtering out low-quality cells, which held true across all cell populations (Fig. S3b,c).

**Figure 3.**
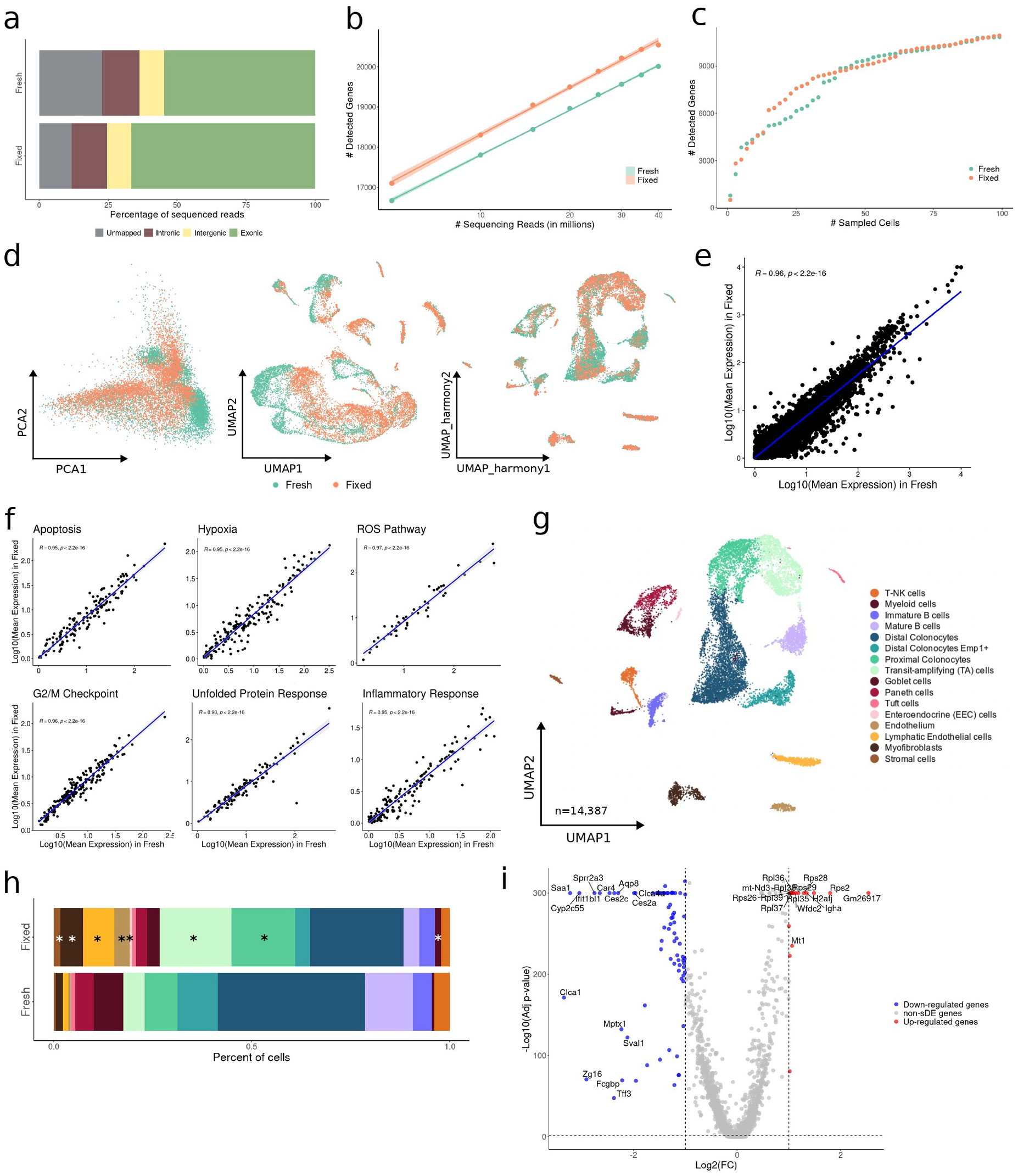
FixNCut protocol tested in mouse colon samples. (**a**) Mapping analysis of sequencing reads within a genomic region. (**b**) Comparative analysis of the number of detected genes based on sequencing reads using a linear model. (**c**) Cumulative gene counts analyzed using randomly sampled cells. (**d**) Principal component analysis (PCA), Uniform Manifold Approximation and Projection (UMAP) prior data integration, and *harmony* integrated UMAP representation of gene expression profile variances of fresh and fixed samples. (**e-f**) Linear regression model comparing average gene expression levels of expressed genes (**e**) and biological hallmarks, including apoptosis, hypoxia, reactive oxygen species (ROS), cell cycle G2/M checkpoint, unfolded protein response (UPR), and inflammatory response genes (**f**). The coefficient of determination (R2) computed with Pearson correlation and the corresponding p-values are indicated. (**g**) UMAP visualization of 14,387 fresh and fixed mouse colon cells, colored by 16 cell populations. (**h**) Comparison of cell population proportions between fresh (n = 6009) and fixed (n = 8378) mouse colon samples with the Bayesian model scCODA. Asterisks (*) indicate credible changes, up-regulated for the fixed sample. (**i**) Differential gene expression analysis between fresh and fixed samples. The top differentially expressed genes (DEGs) with significant adjusted p-values (FDR) < 0.05, up-regulated (red), and down-regulated (blue) with Log2FC > 1 are indicated.

The overlap of HVGs between the fresh and fixed colon samples was slightly lower than that observed in lung tissues (>60%) (Fig. S3d). Further, we identified batch-effects associated with the sample preparation protocol as shown in both PCA and UMAP representations, which were removed by sample integration (Fig. 3d; Fig. S3e). Despite these differences, there was a high correlation in mean gene expression values between the two protocols (R2 = 0.96, p < 2.2e-16), confirming globally conserved expression profiles (Fig. 3e). Notably, cell populations that exhibited a diminished correlation between the fresh and fixed samples coincided with cell types that were specifically enriched in the fixed sample (Fig. S3f).

We captured a total of 14,387 cells that were clustered into 16 cell populations, representing both immune and colon-epithelium cells (Fig. 3g; Fig. S3f). All cell types were detected in both conditions, but we observed a clear shift in cell type composition with an enrichment of sensitive epithelial and stromal cells in the fixed sample (Fig. 3h). Differential expression analysis revealed higher representation of ribosomal protein and mitochondrial genes in the fixed sample, mostly explained by the larger capture of actively cycling epithelial cell population known as transit-amplifying (TA) cells (Fig. 3i; Additional file 5: Table 4). In line, both epithelial and stromal populations were also enriched in mitochondrial and ribosomal protein genes (Additional file 8: Table 7). Additionally, GSEA by cell population showed enrichment of ribosomal-dependent and metabolic processes pathways in fixed cells, especially in the sensitive populations (Additional file 9: Table 8). Together, our results demonstrate the FixNCut protocol to enhance library complexity and quality metrics, while also capturing fragile epithelial and stromal cell populations from both lung and colon tissues. Thus, DSP-based fixation preserves the integrity of tightly connected cell types that are otherwise difficult to isolate for single-cell experiments, resulting in an improved representation of cell types and states in these solid tissues.

#### Long-term storage of fixed tissues

Conducting multi-center clinical studies can be challenging due to centralized data production and the need for storage and shipment. To address this challenge, we evaluated the combination of the FixNCut protocol with cryopreservation (cryo) to allow for the separation of sampling time and location from downstream data generation, while preserving sample composition and gene expression profiles. To test this, fresh mouse lung samples were minced, mixed, and split into three pools for fixation-only, cryo-only, and fixation/cryo sample processing. After single-cell capture and sequencing (10x Genomics, 3’ RNA v3.1), all three libraries showed comparable statistics of mapped and exonic reads across conditions, indicating successful preservation of the transcriptome (Fig. 4a). We also observed a similar relationship between the number of detected genes and the sequencing depth for all three protocols (Fig. 4b), which was consistent considering the detected UMIs and genes across individual cells (Fig. S4a) and when accumulating gene counts across multiple cells (Fig. 4c). After removing low-quality cells and neutrophils, we found highly comparable distributions for the main QC metrics across all samples. However, we noticed a small increase in the percentage of mitochondrial gene expression detected in the fixed/cryo sample (Fig. S4b). Similarly, the different cell populations showed consistent QC across conditions (Fig. S4c).

**Figure 4.**
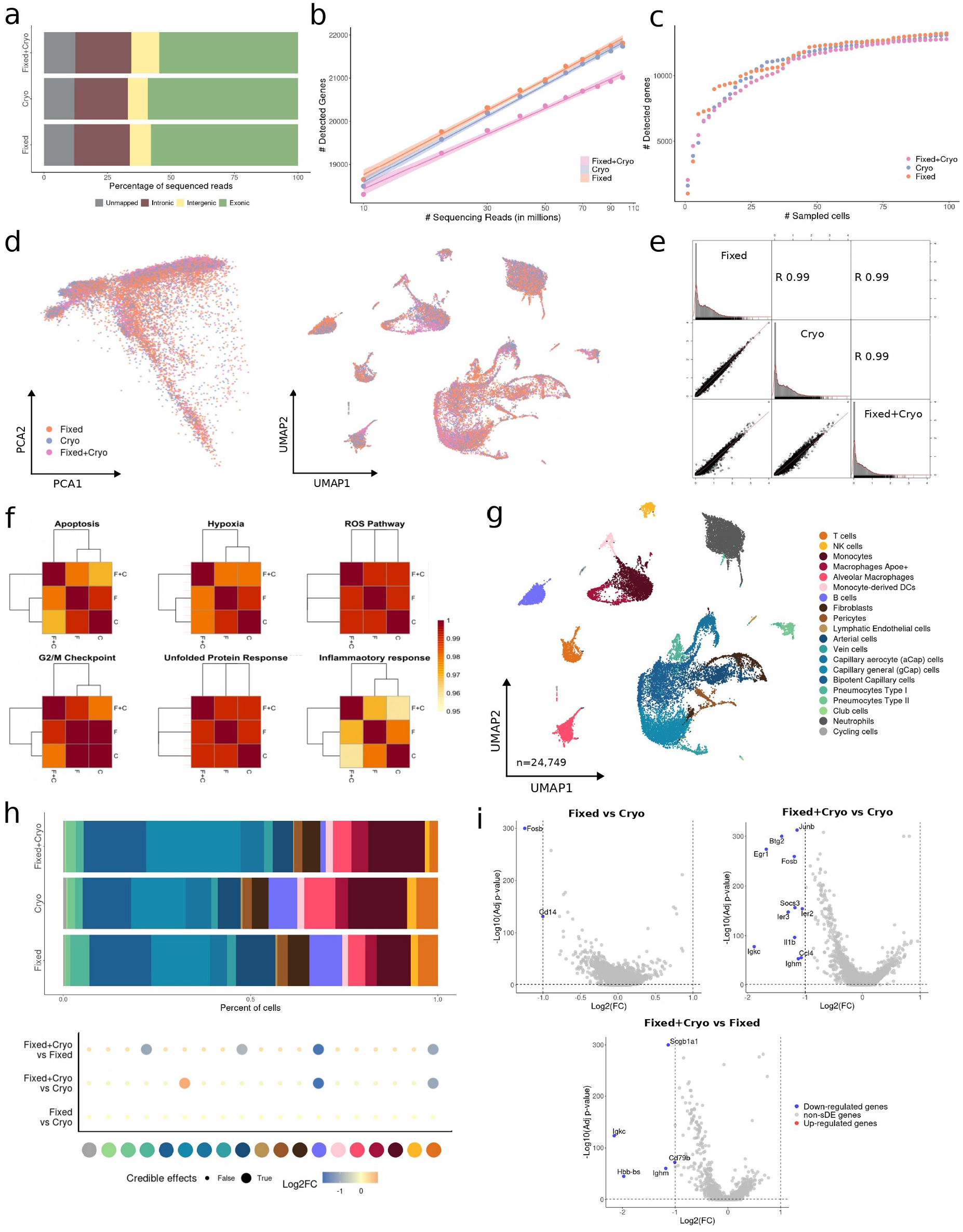
Long-term storage of fixed mouse lung samples. (**a**) Mapping analysis of sequencing reads within a genomic region. (**b**) Comparative analysis of the number of detected genes based on sequencing reads using a linear model. (**c**) Cumulative gene counts analyzed using randomly sampled cells. (**d**) Principal component analysis (PCA) and Uniform Manifold Approximation and Projection (UMAP) representation of gene expression profile variances of fixed, cryopreserved and fixed+cryopreserved samples. (**e**) Linear regression model comparing average gene expression levels of expressed genes across protocols used. The coefficient of determination (R2) computed with Pearson correlation is indicated. (**f**) Hierarchical clustering of coefficient of determination (R2) obtained for all pair comparisons across protocol for biological hallmarks, including apoptosis, hypoxia, reactive oxygen species (ROS), cell cycle G2/M checkpoint, unfolded protein response (UPR), and inflammatory response genes. (**g**) UMAP visualization of 24,749 fixed, cryo, and fixed+cryo mouse lung cells, colored by 20 cell populations. (**h**) Comparison of cell population proportions between fixed (n = 8645), cryopreserved (n=7006), and fixed+cryopreserved cells (n = 4862). The top figure shows the difference in cell population proportions between fixed, cryo and fixed+cryo samples, and the bottom figure shows the results of the compositional cell analysis using the Bayesian model scCODA. Credible changes and Log2FC are indicated. (**i**) Differential gene expression analysis across conditions: fixed vs cryo (**top-left**), fixed+cryo vs cryo (**top-right**), and fixed+cryo vs fixed (**bottom**). The top differentially expressed genes (DEGs) with significant adjusted p-values (FDR) < 0.05, up-regulated (red), and down-regulated (blue) with Log2FC > 1 are indicated.

We confirmed the absence of technical biases after cryopreserving fixed samples, as indicated by a high overlap (>70%) of HVGs across all three protocols (Fig. S4d). In addition, both PCA and UMAP dimensionality reduction plots showed no discernible biases between preservation protocols (Fig. 4d). We also observed highly comparable expression profiles and gene programs when correlating the mean gene expression values for all protocol comparisons (R2 > 0.99, p < 2.2e-16) (Fig. 4e). Moreover, there was no appreciable alteration in biological processes at the gene program or population level when comparing across protocols (Fig. 4f; Fig. S4e).

We next analyzed 24,749 mouse lung cells processed with the three different protocols and annotated 20 lung and tissue-resident immune cell populations (Fig. 4g; Fig. S4f). After removing neutrophils (4,236 cells), all cell types and states were found across the three conditions at similar proportions. However, we observed slight changes in composition, with a decrease of immune cells (B and T Lymphocytes) in the fixed/cryo sample, an increase of gCap compared to the cryo sample, and a subtle decrease in arterial and pneumocyte type I cells compared to the fixed sample (Fig. 4h). Additionally, we observed down-regulation of genes associated with immune function (e.g. Ighm, Igkc, Cd79b, Il1b, Ccl4, Scgb1a1) in the fixed/cryo, explained by the aforementioned shift in cell type composition. Importantly, the cryo-only sample showed up-regulated genes related to stress response, such as Junb or Fosb (Fig. 4i; Additional file 5: Table 4). A closer inspection of the different cell populations validated the expression of stress-related genes across all cells in the cryo-only compared fixed/cryo samples (Additional file 10: Table 9). Accordingly, GSEA detected an enrichment of regulatory or response pathways for almost all cryopreserved cell types compared to fixed/cryo samples (Additional file 11: Table 10). These results support the feasibility of cryopreservation after fixation to combine the robustness and logistical advantages of the respective methods for scRNA-seq experiments.

#### Minimization of technical artifacts in FixNCut tissue samples

Fixing tissues after sample collection preserves the natural state of a cell and avoids technical biases, previously shown to affect bulk and single-cell transcriptomics analysis (2– 4,19). In addition to the above mentioned differences in stress-response genes, we further aimed to demonstrate the ability of FixNCut to preserve gene expression profiles by examining previously identified artifact signatures. Specifically, we investigated condition-specific gene signatures from published studies using our mouse lung and colon data (see Methods).

After performing replicate downsampling on mouse lung samples, we found that fixed samples had significantly lower dissociation/temperature-signature scores compared to fresh samples (p < 2.2e-16), with the first replicate being more affected by external stress (Fig. 5a). We observed that external tissue stressors had a greater impact on fresh lung resident cells compared to the infiltrating immune cell fraction (Fig. 5b). Interestingly, the signature scores for these populations displayed a bimodal-like distribution, indicating an uneven effect within cell populations (Fig. 5b). Additionally, stress-signature genes were differentially expressed in the fresh lung samples (Fig. 5c) and sampling-time signatures, originally defined for PBMCs, showed a significant decrease upon fixation.

**Figure 5.**
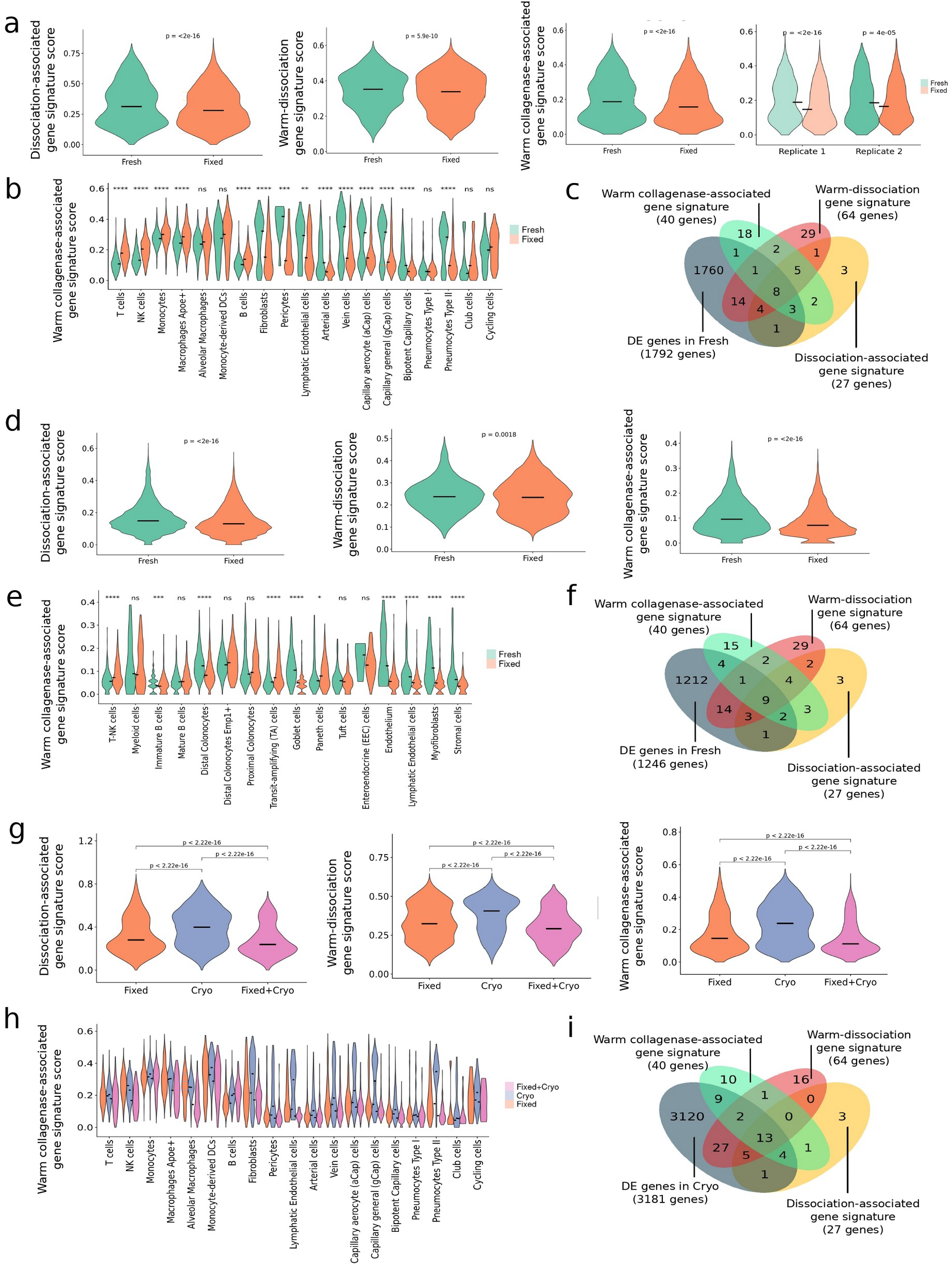
Minimization of technical artifacts using FixNCut protocol on mouse tissues. This figure show the impact of various dissociation-induced gene signature scores, including dissociation-associated on mouse muscle stem cells (19), warm-dissociation on mouse kidney samples (3), and warm collagenase-associated on mouse primary tumors and patient-derived mouse xenografts (2), across mouse tissues and processing protocols used. All statistical analysis between protocols were performed using the Wilcoxon signed-rank test; significance results are indicated with the adjusted p-value, either with real value or approximate result (ns, p>0.05, * p<=0.05, ** p<=0.01, *** p<=0.001, **** p<=0.0001). (**a**) Violin plots of dissociation-induced gene signatures scores for fresh and fixed mouse lung. (**b**) Score of warm collagenase-associated gene signature (2) for fresh and fixed mouse lung samples across cell populations. (**c**) Overlap of differentially expressed genes in the fresh mouse lung samples with genes from the three dissociation-induced signatures. (**d**) Violin plots of dissociation-induced gene signatures scores for fresh and fixed mouse colon. (**e**) Score of warm collagenase-associated gene signature (2) for fresh and fixed mouse colon samples across cell populations. (**f**) Overlap of differentially expressed genes in the fresh mouse colon sample with genes from the three dissociation-induced signatures. (**g**) Violin plots of dissociation-induced gene signatures scores for fixed, cryo and fixed+cryo mouse lung. (**h**) Score of warm collagenase-associated gene signature (2) for fixed, cryo and fixed+cryo mouse lung samples across cell populations. (**i**) Overlap of differentially expressed genes in the cryopreserved mouse lung sample with genes from the three dissociation-induced signatures.

Similarly, fixed colon samples showed a significantly larger decrease in dissociation/temperature-stress signature scores compared to fresh samples (Fig. 5d). Here, we observed a lineage-dependent impact of cell stress; colonocytes were greatly affected with differences in subtypes (more immature cells being more affected), whereas immune cells largely escaped stress biases (Fig. 5e). Secretor, endothelial, and stromal cells suffered the largest dissociation-related stress in the fresh samples, which was drastically reduced upon fixation (Fig. 5e). Moreover, stress-signatures genes were also differentially expressed in the fresh colon sample (Fig. 5f).

Furthermore, we demonstrated the effectiveness of FixNCut for long-term sample storage by examining the dissociation/temperature-stress signature scores in fixed-only, cryo-only, and fixed/cryo mouse lung samples. Our results showed that cryopreserved samples had the highest stress-related signature scores, whereas fixed/cryo samples had the lowest scores for all assessed signatures (p < 2.2e-16) (Fig. 5g). We also observed a reduction in stress signature scores at the cell-type level for fixed-only and fixed/cryo samples compared to cryo-only samples (Fig. 5h). Interestingly, the stress signature score for endothelial and stromal cells exhibited a bimodal distribution exclusively in the cryo-only sample, with cells showing larger dissociation-related effects in the same population (Fig. 5h), consistent with previous observations. Over 70% of signature-specific genes were significantly differentially expressed in the cryo-only sample, an even higher proportion compared to fresh lung and colon samples (Fig. 5i).

#### Moving towards the use of FixNCut on clinical samples

As a proof-of-concept for a multi-center clinical research study focused on autoimmune diseases, we evaluated the performance of FixNCut on human patient biopsies. To this end, we obtained fresh colon biopsies from two IBD patients in remission. The biopsies were mixed and split into four aliquots, which were processed as follows: fresh, fixed-only, cryo-only or fixed/cryo. The fixed human colon samples exhibited a similar proportion of reads mapped to the reference genome and exonic regions as mouse colon tissues (Fig. 6a) and displayed increased library complexity (Fig. 6b) and UMIs at the cell level (Fig. S5a), compared to the fresh and cryo-only samples. The cumulative detected gene count was also highest for the fixed biopsy (Fig. 6c). The main quality control metrics had similar distributions, with slightly improved median UMI, gene counts, and reduced mitochondrial gene percentage for the fixed sample (Fig. S5b), which were consistently observed across almost all populations (Fig. S5c). We observed an overlap (>50%) of HVGs across all conditions (Fig. S5d) and although only subtle batch effects were found, we performed sample integration to ensure a consistent cell annotation across samples (Fig. 6d). Gene expression profiles and gene programs significantly correlated across all samples (R2 > 0.96, p < 2.2e-16) (Fig. 6e,f), with a slightly reduced correlation observed in M0 macrophages and stromal cells (Fig. S5e).

**Figure 6.**
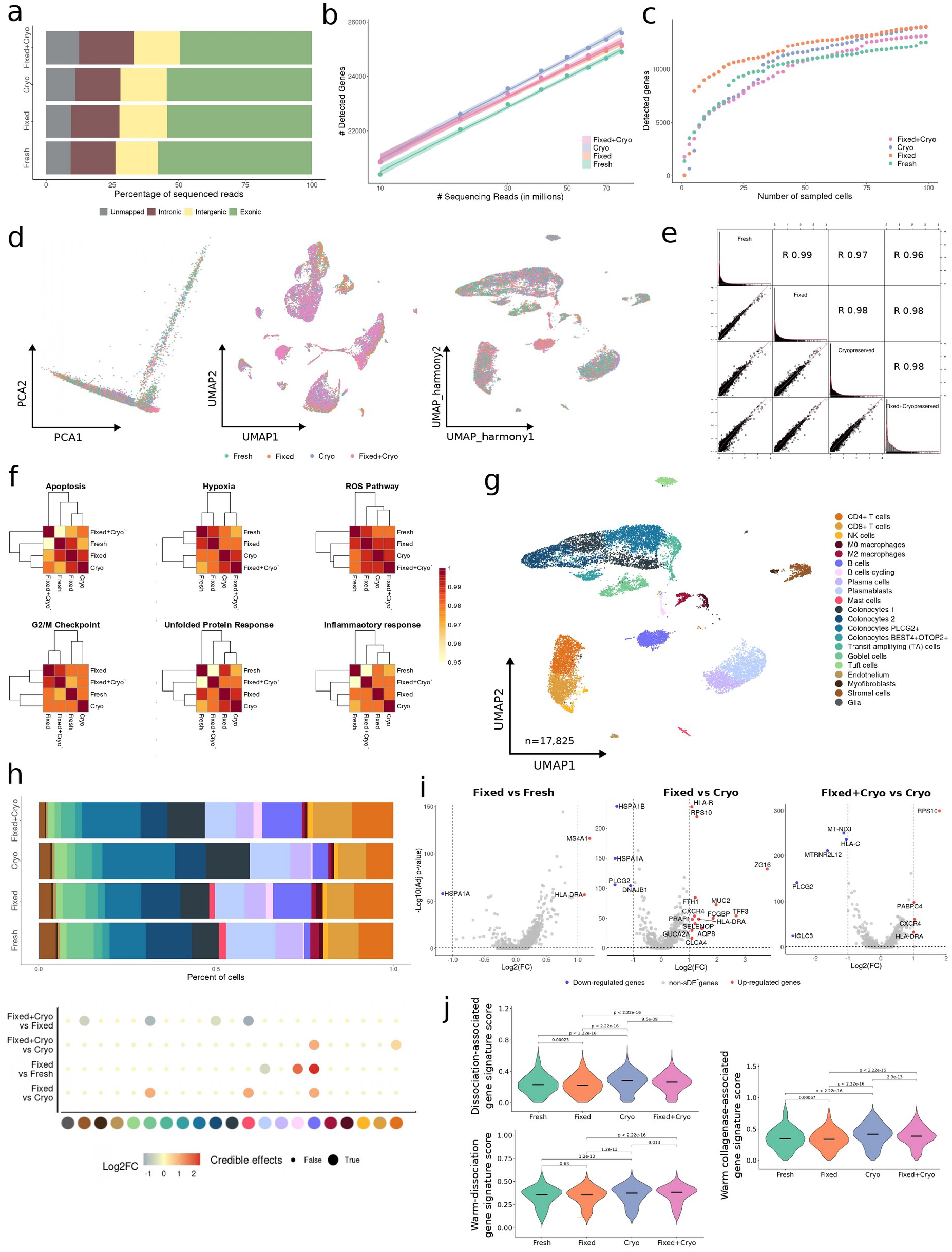
FixNCut protocol tested in human colon biopsies. (**a**) Mapping analysis of sequencing reads within a genomic region. (**b**) Comparative analysis of the number of detected genes based on sequencing reads using a linear model. (**c**) Cumulative gene counts analyzed using randomly sampled cells. (**d**) Principal component analysis (PCA) and Uniform Manifold Approximation and Projection (UMAP) representation of gene expression profile variances prior data integration, and *harmony* integrated UMAP representation of gene expression profile variances of fresh, fixed, cryopreserved and fixed+cryopreserved samples. (**e**) Linear regression model comparing average gene expression levels of expressed genes across protocols used. The coefficient of determination (R2) computed with Pearson correlation is indicated. (**f**) Hierarchical clustering of coefficient of determination (R2) obtained for all pair comparisons across protocols for biological hallmarks, including apoptosis, hypoxia, reactive oxygen species (ROS), cell cycle G2/M checkpoint, unfolded protein response (UPR), and inflammatory response genes. (**g**) UMAP visualization of 17,825 fresh, fixed, cryopreserved, and fixed+cryopreserved human colon cells, colored by 21 cell populations. (**h**) Comparison of cell population proportions between fresh (n = 5759), fixed (n = 4250), cryo (n = 3489), and fixed+cryo (n = 4327) cells. The bottom figure shows the results of compositional cell analysis using the Bayesian model scCODA. Credible changes and Log2FC are indicated. (**i**) Differential gene expression analysis across conditions: fixed vs fresh (**top-left**), fixed vs cryo (**top-right**), fixed+cryo vs cryo (**bottom-left**), and fixed+cryo vs fixed (**bottom-right**). Significant adjusted p-values (FDR) < 0.05, up-regulated (red), and down-regulated (blue) genes with Log2FC > 1 are indicated. The top DE genes are included in the plot. (**j**) Violin plots for stress-related gene signature score (2,3,19) for human colon biopsies across protocols. Statistical analysis between protocols was performed using the Wilcoxon signed-rank test.

By jointly analyzing 17,825 IBD colon cells across protocols, we identified 21 major cell types, including both colon and tissue-resident as well as infiltrated immune cells (Fig. 6g; Fig. S5f). All cell types and states were found across all protocols at similar proportions, although the number of high-quality cells was reduced in the cryo-only sample (Fig. 6h). Notably, the FixNCut protocol captured larger proportions of CD4+ T and B cells compared to fresh or cryo-only samples, among other minor changes (Fig. 6h). We also found that fixed samples had down-regulation of heat-shock proteins (HSP), such as HSPA1A, HSPA1B and DNAJB1, and up-regulation of B cell-specific genes, including MS4A1, HLA-DRA, HLA-B, and CXCR4, when compared to fresh and cryo-only human colon samples (Fig. 6i; Additional file 12: Table 11). In line with the findings from the mouse experiment, we observed that fixed human colon samples exhibited increased expression of ribosomal genes (related to TA cells) at the cell population level, whereas fresh samples showed higher mitochondrial expression. Apart from increased HSP genes in the cryo-only sample, no other significant differentially expressed genes were found between the conditions (Additional file 13: Table 12; Additional file 14: Table 13). GSEA revealed the cryo-only sample to be enriched in stress pathways (response to external stimuli such as stress, temperature, oxidative stress, and protein folding), a pattern also observed in the fixed+cryo sample compared to fixed, but restricted to immune cell types (Additional file 15: Table 14; Additional file 16: Table 15).

By comparing published stress signatures (see Methods) across fresh, fixed-only, cryo-only, and fixed/cryo samples, we found no significant differences between the fixed and fresh samples, whereas the cryo-only sample had significantly higher scores than the fresh sample, with the fixed/cryo sample presenting a lower stress scores than the cryo-only sample; indicating a reduction of gene expression artifacts by fixation for long-term storage in patient biopsies (Fig. 6j). We observed no significant cell population-specific effect, however, stromal cells had the highest score compared to other (Fig. S5g). These findings provide proof-of-concept evidence for the applicability and value of FixNCut in improving the robustness of data generation in the clinical setting.

### Expanding the application of FixNCut to various single-cell techniques

Next, we assessed the compatibility of the FixNCut protocol with single-cell application variants that involve cell labeling with antibodies or lipids targeting the cell membrane (e.g., FACS, CITE-seq, cell hashing). To this end, we stained fresh, cryopreserved and fixed PBMCs and colon tissue samples with fluorescent antibodies or lipid-modified oligos (LMOs), and were analyzed by flow cytometry.

First, we analyzed a cohort of cryopreserved and cryo+fixed human PBMCs (n=3) stained with anti-CD3, CD4, CD8, CD19 monoclonal antibodies (mAbs). The analysis of cell morphology and viability showed that DSP fixation induced slight changes in cell size, internal complexity, and membrane permeability of PBMCs. Specifically, lymphocytes showed a decrease in forward scatter (cell size), while monocytes showed a decrease in side scatter (internal complexity) (Fig. S6a,b).

Next, cells were gated based on the expression of CD3, CD4 and CD8 to characterize all T cell subtypes, and CD19 for B cells. We confirmed that DSP fixation did not alter the percentage of any of the immune cell types analyzed (Fig. 7a). However, using the same amount of antibodies, the mean fluorescence intensity (MFI) was higher in the cryopreserved compared to the cryo+fixed PBMCs (Fig. 7b). To ensure cryopreservation did not introduce any bias, we analyzed another cohort of fresh and fixed human PBMCs (n=4) stained with anti-CD45, CD3, CD4, CD8 mAbs and against ubiquitously expressed human surface proteins (β2M and CD298) and LMOs for cell hashing and multiplexing. DSP fixation led to an increased binding of DAPI, Annexin V (apoptotic cell marker), and propidium iodine (necrotic cell marker) in PBMCs, indicating reduced membrane integrity, particularly after storing cells at 4 °C for 2 days (Fig. S6c). These changes in morphology and membrane integrity should be taken into account when working with mixed study designs including both fixed and fresh samples. Despite observing a minor decrease in the MFI for most of the tested antibodies in fixed samples, we did not detect significant differences in cell type composition comparing fresh to fixed cells (Fig. 7c; Fig. S6d). Similarly, whereas PBMCs labeled with β2M and CD298 antibodies showed no differences in MFI between protocols, cells stained with LMOs revealed minor but noticeable reduction in signal strength in fixed cells (Fig. 7d).

**Figure 7.**
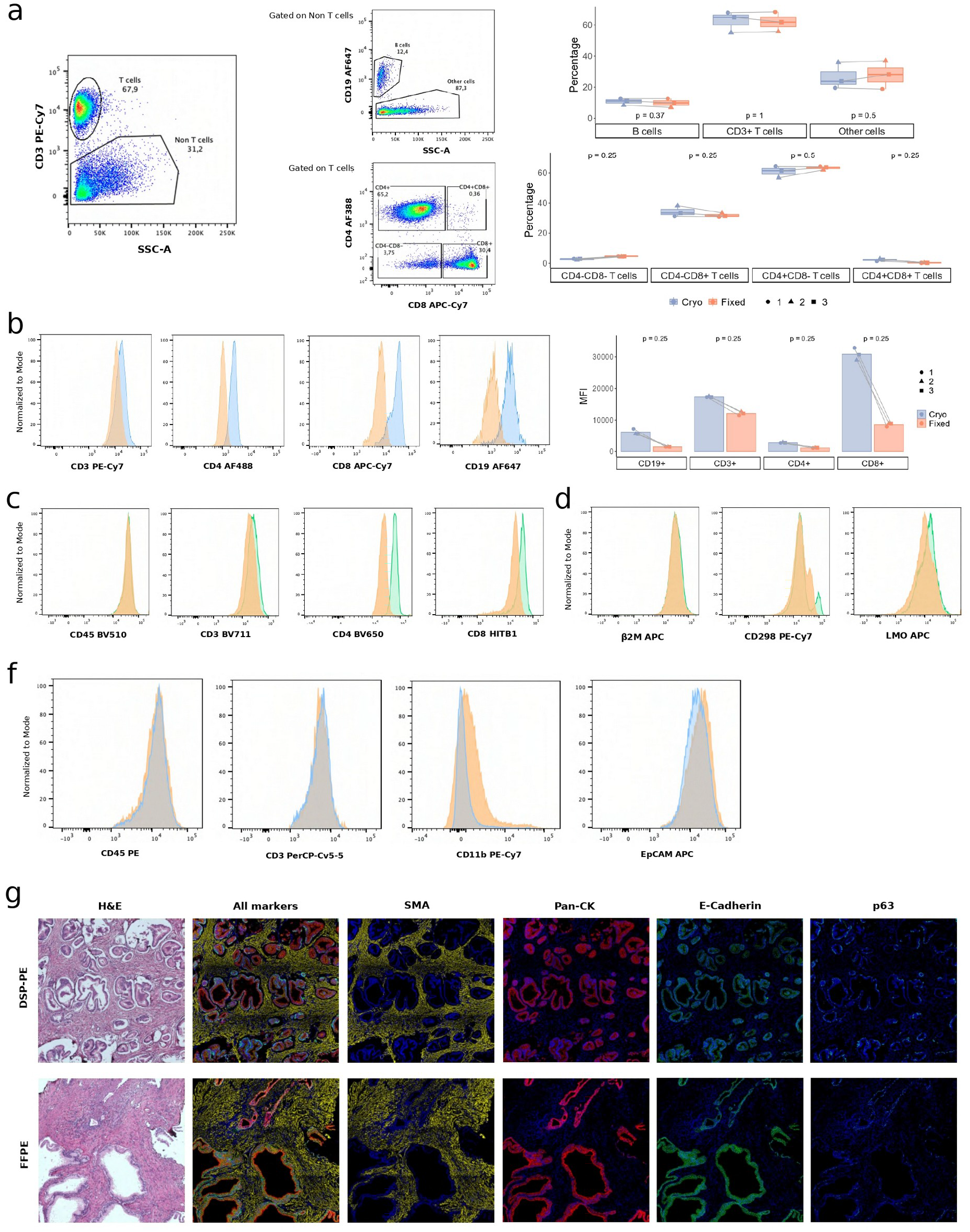
Fluorescent antibody labeling of membrane proteins in fixed cells and tissues in mouse and human. (**a**) Representative gating strategy of one experiment analyzed by flow cytometry with cryopreserved and cryopreserved+fixed PBMCs from healthy donors (n =3). PBMCs were stained with anti-human CD45, CD3, CD19, CD4 and CD8 monoclonal antibodies (mAbs). T cells were selected by the positive expression of CD3, whereas B cells were selected from the CD3 negative fraction (Non T cells) and by the positive expression of CD19. CD4 positive and CD8 positive T cells were selected from CD3 positive T cells. Box plots show the percentage of positive cells in each subpopulation for cryo (blue) and cryo+fixed (orange) PBMCs analyzed by flow cytometry. (**b**) Representative histograms of the mean fluorescent intensity (MFI) with cryopreserved and cryopreserved+fixed PBMCs from healthy donors (n=3) stained with anti-human CD45, CD3, CD19, CD4 and CD8 mAbs. Bar plots show the MFI expression for three fresh and fixed PBMC samples analyzed by flow cytometry. (**c-d**) Representative histograms of MFI of fresh and fixed PBMCs from healthy donors (n=4) stained with anti-human CD45, CD3, CD4 and CD8 mAbs, anti-β2M, anti-CD298, and LMOs analyzed by flow cytometry. (**f**) Representative histograms of the MFI from cryopreserved (blue) and cryopreserved+fixed (orange) human colon samples processed and stained with anti-human CD45, CD3, CD11b, and EpCAM mAbs and analyzed by flow cytometry. (**g**) Multiplex fluorescence tissue imaging of a human prostate cancer section, DSP-fixed (**top**) or formalin-fixed (**bottom**) paraffin-embedded, captured using Phenocycler. Images show hematoxylin and eosin staining, five-color overlay, and individual SMA, Pan-CK, E-cadherin, and p63 antibody staining.

Similarly, staining dissociated human colon biopsies with anti-CD45, CD3, CD11b and EpCAM antibodies showed similar MFI in cryopreserved and fixed cells (Fig. 7f; Fig. S6e). Our results showed that DSP fixation is compatible with cell labeling of cells with antibodies. Nevertheless, we recommend the optimization of labeling conditions and flow cytometry protocol of fixed samples, depending on antibody sensitivity, antigen abundance, and downstream applications, to further improve results.

Lastly, we have effectively conducted multiplexed immunofluorescence tissue imaging on DSP-fixed paraffin-embedded prostate cancer and its formalin-fixed paraffin-embedded counterpart, the latter considered as ‘gold-standard’ (Fig. 7g). This demonstrates the feasibility of DSP fixation in spatial-omics, particularly for spatial proteomics using technologies like Phenocycler (Akoya Biosciences).

## Discussion

In this study, we introduced the FixNCut protocol, a novel approach that combines sample fixation with subsequent tissue dissociation to overcome several limitations in generating single-cell data. While DSP fixation has previously been used to fix K562 cells and keratinocytes prior to sequencing (15,16), or in combination with single-cell technologies to explore the adaptive immune repertoire (17), our study is the first to utilize the reversible properties of the DSP fixative with standard enzymatic dissociation on solid biopsies in the context of single-cell transcriptomic studies.

The FixNCut protocol offers significant advantages, including the ability to fix tissue prior to digestion, providing a snapshot of the cell transcriptome at sampling time and minimizing technical artifacts during tissue processing. As the reversible fixative targets proteins rather than nucleic acids, our approach ensures RNA integrity, library complexity, cellular composition, and gene expression comparable to those of the gold standard fresh RNA sequencing assays. With FixNCut, time and location constraints for sample collection and processing are removed, making it an ideal strategy for research studies involving multiple groups, institutions, and hospitals worldwide.

Standardizing protocols for clinical biopsy collection and downstream processing poses a significant challenge. While single-cell profiling technologies have proven to be highly useful and straightforward for PBMCs and other cells loosely retained in secondary lymphoid tissues, the effective isolation of single cells from solid tissues, such as tumors, remains a technical hurdle. In such cases, single cells may be tightly bound to extracellular scaffolds and neighboring cells, making dissociation and isolation difficult (20). Additionally, preserving the single-cell transcriptome before scRNA-seq is crucial, particularly when processing multiple samples from biological replicates simultaneously, as it can reduce the need for immediate time-consuming single-cell isolation protocols, such as dissociation, antibody staining or FACS-based isolation (20). Recent studies have highlighted the impact of collection and dissociation protocols on cell type proportions and transcriptome profiles in multiple tissue contexts (2–4). Furthermore, dissociation-related effects have also been observed in cryopreserved human gut mucosa samples (21) and renal biopsies (22). To overcome these issues, a one-step collagenase protocol was used for intestinal biopsies when no cell type enrichment was required (21). Meanwhile, the use of cold-active proteases on kidney samples resulted in fewer artifacts, but inefficient tissue dissociation (22). In addition, studies have shown that neural cell populations (NCPs) suspension without methanol preservation also experiences alterations in cellular composition and gene expression (23). For fragile tissue biopsies, such as pancreas or skin, which contain delicate cell populations, cryopreservation and cellular dissociation steps may introduce biases on the cellular composition. While this manuscript was being prepared, the preprint by Fortmann *et al.* was made available in BioRxiv (24). We had previously conducted internal testing and found the external use of VivoFix to be challenging, and it did not yield successful results for us. However, we acknowledge the potential utility of this protocol in the hands of the authors.

To address these challenges, we developed FixNCut, an approach that involves reversible fixation of the tissue at the time of collection to prevent further transcriptional changes during downstream processing. This is followed by standard dissociation and storage procedures. At the core of our approach is the use of Lomant’s reagent/DSP, a reversible fixative that can easily penetrate cell membranes and preserve tissue characteristics. In this study, we compared fresh and fixed lung and colon samples from different species and experimental scenarios, and found comparable results. Additionally, we demonstrated the versatility of the FixNCut for long-term storage by cryopreservation following sample fixation, making it a suitable protocol for its use in more complex and challenging research scenarios. While the lung represents a more resilient tissue for sample processing, without the introduction of major changes in gene expression or cellular composition, colon tissue is very sensitive. Here, we demonstrated decreased RNA quality and a shift in cellular composition in fresh compared to fixed samples, even under standard experimental conditions. We predict that under more stressful conditions, such as therapeutic intervention models, these biases will be even more pronounced.

In single-cell analysis, antibodies are commonly used to select cells of interest by standard FACS enrichment or to quantify cell surface proteins using sequencing (e.g. CITE-Seq). Unlike other fixative agents, such as methanol, which induces protein unfolding and precipitation, or formalin, which non-selectively cross-links proteins, DNA and RNA (reducing immunoreactivity with target-specific antibodies), the FixNCut protocol has the advantage that cells can be readily stained with antibodies and LMOs. This allows cell labeling or hashing before single-cell analysis. Hence, we assume that this protocol is adaptable and compatible with multiple single-cell modalities, including but not limited to CITE-seq, as well as other droplet or microwell platforms. This versatility makes it a powerful tool for designing flexible and robust studies, being applicable to different tissue, species, or disease conditions.

The FixNCut protocol offers a straightforward way to preserve biopsies for various research contexts, including animal models at research institutes and patient biopsies collected at hospitals. We have demonstrated that FixNCut can be applied in clinical settings, where samples are collected at separate locations and time from their downstream processing steps. The FixNCut protocol’s versatility and compatibility with various 10x Genomics applications, such as CITE-Seq, ATAC, and Multiome, in addition to sc/snRNA-seq, make it a promising tool for preserving samples in different areas (e.g., oncology and autoimmunity).

### Conclusions

We demonstrate the FixNCut protocol to preserve the transcriptional profile of single cells and the cellular composition of complex tissues. The protocol enables the disconnection of sampling time and location from subsequent processing sites, particularly important in clinical settings. The protocol further prevents sample processing artifacts by stabilizing cellular transcriptomes, enabling robust and flexible study designs for collaborative research and personalized medicine.

## Methods

### Sample collection

#### Human PBMCs isolation

Peripheral venous blood samples were collected from voluntary blood donors using ACD-tubes and stored at 4°C. PBMCs isolation was performed using Ficoll density gradient centrifugation. Briefly, 10 mL of blood were diluted with an equal volume of 1X PBS and carefully layered onto 15 mL of Lymphoprep (PN. 15257179, STEMCELL Technologies) followed by centrifugation for 20 min at 800 x g and room temperature (RT) (with acceleration and brake off). After centrifugation, PBMCs were collected with a sterile Pasteur pipette, transferred to a 15 mL tube, and washed twice with 10 mL of 1X PBS by centrifugation for 5 min at 500 x g at RT. PBMCs were resuspended in 1X PBS + 0.05% BSA, and cell number and viability were measured with LUNA-FL^TM^ Dual Fluorescence Cell Counter (LogosBiosystem).

#### Mouse lungs collection

C57BL/6 mice were purchased from Janvier Laboratories at 6 weeks of age and sacrificed between week 7 and 9 by CO_2_ asphyxiation. Lung samples of replicate 1 were not subjected to perfusion, whereas lungs of replicate 2 were perfused prior to collection. To perfuse the lungs, a 26G syringe was used to inject 3 mL of cold Hank’s Balanced Salt Solution (HBSS) into the right ventricle of the heart, which resulted in the lungs turning white after injection. Mice from both experimental replicates were then carefully dissected for further processing.

#### Mouse colon collection

Mice were sacrificed using as described above, and the colon was collected and washed with HBSS using a syringe to remove feces. The collected colon samples were transported from the facility to the lab in complete DMEM medium on ice. Upon arrival, the samples were extensively washed with ice-cold PBS and then cut into 3×3 mm pieces on a Petri dish using a sterile razor blade. The tissue pieces were then fixed as previously described.

#### Human colon biopsies

Colonic biopsies were collected from an ulcerative colitis patient in remission and placed in HBSS (Gibco, MA, USA) until processing, which was completed within an hour. The biopsies were split into four different conditions: fresh, fixed, cryopreserved and fixed+cryopreserved. For fixation, the biopsies were treated as previously described for mouse lung tissue. Cryopreservation of fresh or fixed biopsies was done by transferring them into 1 mL of freezing media (90% FBS + 10% DMSO, Thermo Scientific) and storing them at -80°C in a Mr. Frosty™ Freezing Container to ensure gradual freezing.

#### Human prostate tissue

Human prostate tissue was collected with informed consent from a 75-year-old patient who underwent a radical prostatectomy procedure for prostate cancer.

### Sample preparation: fixation and cryopreservation

#### Preparation of DSP fixation buffer

A 50X stock solution of DSP (50 mg/mL) was prepared in 100% anhydrous DMSO and stored at -80°C. Prior to use, 10 μL of the 50X DSP was added dropwise to 490 μL of RT PBS in a 1.5 mL tube while vortexing to prevent DSP precipitation. This working solution (1X DSP fixation buffer) was then filtered through a 40 μm Flowmi Cell Strainer (PN. BAH136800040-50EA, Sigma-Aldrich). Table 2 provides detailed instructions for using the FixNCut protocol with both the DSP stock solution and working dilution.

#### PBMCs fixation

1 million cells were split into two separate 1.5 mL tubes, with one tube used fresh (as a non-fixed control sample) while the other was subjected to cell fixation. For fixation, cells were centrifuged at 500 x g for 5 minutes at 4°C, and the resulting pellet was resuspended with 500 μL of 1X DSP fixation buffer and incubated at RT. After 15 minutes, the cells were mixed by pipetting and incubated for an additional 15 minutes. Fixation was stopped by adding 10 μL of 1 M Tris-HCl pH 7.4, and the sample was briefly vortexed and incubated at RT for 5 minutes. Both fresh and fixed samples were centrifuged for 5 minutes at 500 x g at 4°C, and contaminating erythrocytes were eliminated by resuspending the pellets and incubating at RT for 5 minutes with 1X Red Blood Cell lysis solution (PN. 130-094-183, Miltenyi Biotec). Both samples, fresh and fixed, were centrifuged for 5 min at 500 x g at 4⁰C and contaminating erythrocytes were eliminated by resuspending the pellets in 500 μL of PBS and incubating at RT for 5 min upon addition of 10 times volume of 1X Red Blood Cell lysis solution (PN. 130-094-183, Miltenyi Biotec). Cells were then resuspended in an appropriate volume of 1X PBS + 0.05% BSA in order to reach optimal concentration for cell encapsulation (700-1000 cells/μL) and filtered using a pluriStrainer Mini 40 µm (PN. 43-10040-70 Cell Strainer). Cell concentration was verified with LUNA-FL^TM^ Dual Fluorescence Cell Counter (LogosBiosystem).

### Single-cell RNA-seq experimental design (scRNA-seq)

#### Human PBMC 3’ scRNA-seq

Cells from both fresh and fixed PBMCs were processed for single-cell RNA sequencing using the Chromium Controller system (10X Genomics), with a target recovery of 8000 total cells. The Next GEM Single Cell 3’ Reagent Kits v3.1 (PN-1000268, 10X Genomics) were used to prepare cDNA sequencing libraries, following the manufacturer’s instructions. Briefly, after GEM-RT clean up, cDNA was amplified for 11 cycles and then subjected to quality control and quantification using an Agilent Bioanalyzer High Sensitivity chip (Agilent Technologies). The Dual Index Kit TT Set A (PN-1000215, 10X Genomics) was used for indexing cDNA libraries by PCR. The size distribution and concentration of 3’ cDNA libraries were verified again on an Agilent Bioanalyzer High Sensitivity chip (Agilent Technologies). The cDNA libraries were sequenced on an Illumina NovaSeq 6000 using the following sequencing conditions: 28 bp (Read 1) + 8 bp (i7 index) + 0 bp (i5 index) + 89 bp (Read 2) to generate approximately 40,000 read pairs per cell.

#### Mouse lung cryopreservation, fixation and cryopreservation upon fixation

Mouse lungs were harvested and transferred into ice cold complete DMEM medium. Samples were extensively washed with ice-cold PBS, transferred into a Petri dish, and cut into ∼3×3 mm pieces using a razor blade. Tissue pieces were divided into four tubes for each condition. Tissue pieces in tube 1 were cryopreserved in freezing media (50% DMEM + 40% FBS + 10% DMSO) and placed into a Mr. Frosty™ Freezing Container and transferred to a -80°C freezer to ensure a gradual freezing process. The tissue pieces in tubes 2 and 3 were minced with a razor blade (∼1×1 mm) on ice and fixed by submerging them in 500 μL of 1X DSP fixation buffer (freshly prepared, within 5 minutes of use) and incubating at RT for 30 min. After incubation, 10 μL of 1 M Tris-HCl pH 7.5 were added to stop the fixation, and the samples were vortexed for 2-3 seconds and incubated at RT for 5 minutes. After a brief centrifugation, the supernatant was removed, and the tissue pieces were washed once with 1 mL of PBS. Tissue pieces in tube 2 were stored at 4°C in PBS supplemented with 2 U/μL of RNAse inhibitor (Cat. N. 3335402001, Sigma Aldrich) until the following day, while tissue pieces in tube 3 were cryopreserved by adding 10% DMSO to the PBS, transferring the cell suspension into a cryotube, which was placed into a Mr. Frosty™ Freezing Container, and stored at -80°C. The tube containing fresh tissue was stored in complete DMEM on ice and washed again with ice-cold PBS before tissue dissociation.

#### Mouse lung dissociation and scRNA-seq

The day after the collection and storage of samples, the cryopreserved and fixed+cryopreserved samples were quickly thawed in a 37°C water bath and washed with PBS. Similarly, the fixed sample was washed with PBS before tissue dissociation. The samples were then transferred to a Petri dish on ice and minced using a razor blade. Next, the minced samples were incubated in 1 mL of digestion media (200 μg/mL Liberase TL, 100 μg/mL DNase I in HBSS with Ca^2+^Mg^2+^) at 37°C with shaking at 800 rpm. After 15 minutes of incubation, the samples were mixed by pipetting, followed by another 15 minutes of incubation. The cells were then filtered using a pluriStrainer Mini 70 µm (PN. 43-10070-40 Cell Strainer), and the strainer was washed with 10 mL of cold 1X HBSS. The samples were centrifuged at 500 x g for 5 minutes at 4°C, and the cell pellets were resuspended in 100 μL of PBS + 0.05% BSA. Contaminating erythrocytes were lysed using the previously described method. The cells were washed once with 10 mL PBS + 0.05% BSA, resuspended in an appropriate volume of the same buffer, and filtered using 40 μm strainers. The total number of cells was determined using the LUNA-FL^TM^ Dual Fluorescence Cell Counter (LogosBiosystem). The cell concentration of each sample was adjusted to 700-1000 cells/μL and 7000-10.000 cells were loaded into a 10X Chromium controller. Single-cell RNA-sequencing was performed as described above.

#### Mouse colon dissociation

Fresh and fixed colon samples were incubated in 1 mL of digestion media (200 U/mL Collagenase IV, 100 μg/mL DNase I in HBSS w/Ca^2+^Mg^2+^) at 37°C, shaking at 800 rpm for 30 minutes. Samples were mixed by pipetting every 10 minutes during incubation. After the incubation, samples were filtered through a pluriStrainer Mini 70 µm (PN. 43-10070-40 Cell Strainer) and the strainer was washed with 10 mL of cold 1X HBSS. The samples were then centrifuged at 500 x g for 5 min at 4°C and cell pellets were washed twice with cold PBS + 0.05% BSA. Finally, the cell pellets were resuspended in an appropriate volume of the same buffer and filtered using 40 μm strainers. The total cell number was determined with the LUNA-FL^TM^ Dual Fluorescence Cell Counter (LogosBiosystem). The cell concentration of each sample was adjusted to 700-1000 cells/μL and 7000 cells were loaded onto a 10X Chromium controller. Single cell RNA-seq was performed as described above.

#### Human colon biopsies dissociation

Digestion of biopsies to single cell suspension was achieved through mechanical and enzymatic digestion as previously described by Veny *et al*. (24). Briefly, biopsies were washed in complete medium (CM) (RPMI 1640 medium (Lonza, MD, USA) supplemented with 10% FBS (Biosera, France), 100 U/mL penicillin, 100 U/mL streptomycin and 250 ng/mL amphotericin B (Lonza), 10 µg/mL gentamicin sulfate (Lonza) and 1.5 mM HEPES (Lonza)) before being minced and incubated in 500 µL of digestion media (CM supplemented with Liberase TM (0.5 Wünsch U/mL) (Roche, Spain) + DNase I (10 μg/mL) (Roche, Spain)) with agitation (250 rpm) for 1 hour at 37°C. After incubation, the cell suspension was filtered using 50 µm and 30 µm cell strainers (CellTrics, Sysmex, USA) to remove cell aggregates and debris. Cell viability and concentration were determined with the LUNA-FL^TM^ Dual Fluorescence Cell Counter (LogosBiosystem). Approximately 7000 cells were loaded onto the Chromium controller (10x Genomics, CA, USA).

### Flow cytometric analysis

#### Flow cytometry analysis of human PBMCs

Anti-human CD3, CD4, CD8 and CD19 antibodies were tested as follows: Cryopreserved PBMCs obtained from three healthy donors were rapidly thawed in a 37°C water bath. Thawed samples were washed in pre-warmed RPMI media supplemented with 10% FBS (Thermo Fisher Scientific) and centrifuged at 500 x g for 5 min at RT. The supernatant was discarded, and pellets were washed in 10 mL of 1X PBS + 0.05% BSA, centrifuged at 500 x g for 5 min at 4°C, and resuspended in 1 mL of PBS + 0.05% BSA. The cell suspension was then filtered through a 40 µm cell strainer. Cell viability and concentration were verified using a LUNA-FL^TM^ Dual Fluorescence Cell Counter (LogosBiosystem). Each sample was split into two separate tubes, and half of the cells were fixed with 1X DSP fixation buffer as previously described. After fixation, cells were washed and resuspended in 100 μL of Cell Staining Buffer (PN-420201, Biolegend) and stained with 5 μL of each of the four following primary antibodies: anti-human CD3, CD4, CD8, CD19 antibody for 15 min at RT in the dark. Detailed information on the antibodies and reagents used in this study is provided in Table 3. Samples were washed twice with Cell Staining Buffer and resuspended in 0.5-1 mL of PBS + 0.05% BSA. 10 μg/mL DAPI (PN-564907, BD Bioscience) was added to determine cell viability before flow cytometric analysis using the BD FACS Melody Automated Cell Sorter (BD Bioscience) and the BD FACSChorus™ Software. Post-acquisition analysis was performed using FlowJo version 10 (FlowJo LLC).

Anti-human CD45, CD3, CD4, CD8 antibodies and anti-CD298 and β2-microglobulin antibodies were tested as follows: Human PBMCs were isolated from normal donor human buffy coats provided by the Australian Red Cross Blood Service by Ficoll-Paque® density gradient centrifugation. Fresh and fixed PBMCs were incubated with Human BD Fc Block for 10 min at 4°C and then stained for cell surface markers for 30 min at 4°C according to the manufacturer’s recommendation. Annexin V-FITC and PI staining were used to determine viability. Detailed information on the antibodies and reagents used in this study is provided in Table 3. Acquisition was performed on LSR Fortessa X 20 (BD) and analyzed via FlowJo software (FlowJo LLC).

#### LMO sample preparation

The labelling of PBMCs was performed following the protocol previously described by McGinnis *et al.* (25). Briefly, 5 x 105 fresh and fixed PBMCs were washed twice with PBS and labelled with a 1:1 molar ratio of anchor LMO and barcode oligonucleotide for 5 min on ice. Subsequently, both samples were incubated with a co-anchor and Alexa 647 fluorescent oligo feature barcodes at concentrations of 200 nM and 400 nM, respectively, for another 5 min on ice. The cells were then washed twice with ice-cold 1% BSA in PBS. Acquisition was performed on the LSR Fortessa X 20 (BD) and analysis was carried out using FlowJo software (FlowJo LLC). Detailed information on the LMOs used in this study is provided in Table 4.

#### Flow cytometry analysis of human colon biopsy

The single-cell suspensions obtained after biopsy digestion were labelled with the following antibodies: anti-CD45, anti-CD3, anti-CD11b, and anti-EPCAM, according to the manufacturer’s instructions. Detailed information on the antibodies used in this study can be found in Table 3. Cell viability was assessed using the Zombie Aqua Fixable Viability Kit (BioLegend). The cells were then fixed with the Stabilizing Fixative (BD) before being analyzed using the FACSCanto II flow cytometer (BD).

### Multiplexing fluorescence tissue imaging

#### Tissue preparation

The 2 cm x 10 mm tissue sample was divided into two equal halves lengthwise. One half was fixed in 2 ml of 10% neutral buffered formalin (NBF), while the other half was placed in 500 µL of 1X DSP fixation buffer. The NBF-fixed sample was incubated at RT for 4 hours, stored overnight at 4°C, washed 3 times with 1 mL of milliQ water, and then stored in 1 mL of 70% ethanol at 4°C. The DSP-fixed sample was treated with freshly-made DSP fixation buffer, which was replaced every 60 minutes for 4 hours. The fixed sample was then neutralized with 10 µL of 1M Tris-HCl pH 7.4 for 15 minutes at RT, washed 3 times with 1 mL of milliQ water, and placed in 1 mL of 70% ethanol at 4°C. Both NBF- and DSP-fixed samples were embedded in paraffin overnight. Five-micrometer-thick sections were cut from both the formalin-fixed, paraffin-embedded (FFPE) and the DSP-fixed, paraffin-embedded (DSP-PE) tissues, and mounted onto a single poly-L-lysine coated coverslip (22×22 mm, #1.5, Akoya Bioscience #7000005).

#### Antibody staining

The coverslip mounted section was baked at 60°C on a heat block for 1 h to remove paraffin, then deparaffinized in 1X Histo-Choice clearing agent (ProSciTech #EMS64110-01) and rehydrated in ethanol before washing in milliQ water. Antigen retrieval was performed in a pressure cooker on the highest setting for 20 min in 1X citrate buffer, pH 6 (Sigma, #C9999-1000ML). The tissue was blocked using buffers from the commercially available Phenocycler staining kit (Akoya Bioscience, # 7000008) and stained with a 7-antibody panel at RT for 3 h. Detailed information on the antibodies used in this study can be found in Table 5. The antibodies were used at a dilution of 0.9:200 for commercially available Akoya antibody-oligo conjugates and 3.7:200 for antibodies custom-conjugated by Akoya Bioscience (Spatial Tissue Exploration Program (STEP)). After staining, the coverslip was subjected to a post-fixation step. DAPI staining was used to visualize cell nuclei and locate regions of interest on each tissue sample with the Zeiss Axio Observer 7 fluorescent inverted microscope.

#### Phenocycler image acquisition

The Phenocycler microfluidic microscope stage was programmed to acquire two 3 x 3 tiled regions on each tissue using a 20X objective lens, with each tile consisting of a 7-image Z-stack illuminated by LED light to specifically excite either DAPI (for 10 ms in all cycles) or one of three fluorescently labeled reporter oligos (Cy3, Cy5, and Cy7). The software was set to acquire images over 5 cycles, with each cycle consisting of the addition of a set of reporter oligos complementary to the antibody-oligo conjugates detailed in Table 5. During the first and last cycles, no reporter oligos were added to allow for background fluorescence subtraction. The exposure times for each antibody are also provided in Table 5. After imaging was completed, the sample was manually stained with Hematoxylin and Eosin (H&E) following the UofA histology protocol, and bright-field images were captured using the Zeiss Axio Observer 5 fluorescent inverted microscope.

#### Image processing

The acquired images were processed using the Phenocycler CODEX Processor software (Akoya version 1.8.3.14) to deconvolve and stitch them together, resulting in a set of multi-channel QPTIFF files for each region. The levels for each channel were adjusted using QuPath 0.4.0 and the final images were saved as RGB tiff files. However, some antibodies did not produce sufficient signal or acceptable images after processing and were therefore excluded from further analysis.

### Data processing

To profile the cellular transcriptome, we processed the sequencing reads using the CellRanger software package (version 6.1.1) from 10X Genomics Inc. We mapped the reads against either the mouse mm10 or the human GRCh38 reference genome, depending on the samples. In order to avoid any artifacts on the downstream analysis due to differences in sequencing depth among samples, we normalized libraries for effective sequencing depth using “*cellranger aggr*”. This subsampling approach ensures that all libraries have the same average number of filtered reads mapped confidently to the transcriptome per cell.

### Data analysis

All analyses presented in this manuscript were conducted using R version 4.0.5, along with specific analysis and data visualization packages. For scRNA-seq analysis, we used Seurat R package (version 4.0.0) (26), SeuratObject package (version 4.0.1), and other packages specified in the subsequent sections.

#### Assessing quality control parameters of scRNA-seq libraries

To compare the library complexity (total captured genes) across libraries, we investigated the relationship between the cumulative number of detected genes and the library sequencing depth. To achieve this, we downsampled the library sequenced reads assigned to a barcode (excluding background and noisy reads) using the DropletUtils package function “*downsampleReads*”. We utilized various depths for downsampling (steps of 5M or 10M reads, depending on the library quality), which emulates differences in sequencing depth per cell. Moreover, we assessed the distribution of cell complexity (total captured genes and UMIs) per cell sequencing depth across libraries. For both scenarios, we computed a linear model to compare the slope of the regression line for each different library. Ultimately, we computed the cumulative number of detected genes over multiple cells by averaging the total genes after 100 permutations of an increasing number of randomly sampled cells (from 1 to 100, using steps of 2), after running the “*cellranger aggr*” step described above.

#### scRNA-seq quality control and cell annotation

After ensuring that there were no remarkable differences on the main quality control (QC) metrics (library size, library complexity, percentage of mitochondrial and ribosomal expression) among the different samples, we performed an independent QC, normalization and analysis for the libraries from different species and tissue, following the guidelines provided by Luecken *et al.* (27). We removed low-quality cells by filtering out barcodes with a very low number of UMIs and genes, or with a high percentage of mitochondrial expression, as it is indicative of lysed cells. Additionally, we considered removing barcodes with a large library size and complexity. We eliminated genes that were detected in very few cells. Notably, due to the inherent characteristics of colon biopsies (a higher number of epithelial cells, which are less resistant to sample processing), we followed a slightly different QC approach for mouse and human colon samples. In brief, we performed a first permissive QC filtering out cells with very high MT% (>75% for mouse and >85% for human) before proceeding to downstream analysis. We annotated cells to distinguish between the epithelial and non-epithelial fraction. Then, we repeated the QC step, using different thresholds for the epithelial fraction (>60% MT) and for the non-epithelial cells (>50% for mouse and >25% for human). Finally, data was normalized and log transformed.

To achieve successful cell-type annotation combining data from the same tissue and species (mouse and human colon samples), we removed the batch-effect with the *Harmony* integration method (28) using the library as a confounder variable. After integration, we created a k-nearest neighbors (KNN) graph with the “*FindNeighbors*” function using the first 20 Principal Components (PC), followed by the cell clustering with the Louvain clustering algorithm using the “*FindClusters*” function at different resolutions. To visualize our data in a two-dimensional embedding, we run the Uniform Manifold Approximation and Projection (UMAP) algorithm. Then, we performed a Differential Expression Analysis (DEA) for all clusters to determine their marker genes using the normalized RNA counts. To annotate the clusters into specific cell types, we examined the expression of canonical gene markers, compared the results of the DE analysis and referred to gene markers from published annotated datasets. We used the following datasets as references: human PBMCs based on Stuart *et al.* (29), mouse lung based on Angelidis *et al.* (30) and Zhang *et al.* (31), mouse colon based on Tsang *et al.* (32), and human colon based on Garrido-Trigo *et al*. (33). Furthermore, we performed specific cell-type sub-clustering when required a fine-grained resolution to capture a specific cell-state of interest. Doublets and low-quality cells were automatically removed at this point.

### Comparison of gene expression profiles

#### Gene expression correlation analysis

Overall similarity of pseudo-bulk gene expression profiles across experimental conditions was explored using the Pearson correlation (r^2^). To generate the pseudo-bulk profiles, we computed the mean of the log-average gene counts across all cells per sample, as well as for each defined cell population independently. Due to the sparsity of scRNA-seq data, we only computed correlation for the cell-types with over 100 cells and only if over 20 cells were present by condition to compare. Additionally, to avoid biases driven by cell populations with different sizes, we randomly downsampled the number of cells to the condition having fewer cells. To assess the strength and significance of the correlation, we used linear regression models; we considered strong significant linear correlation when r^2^ > 0.9 and p-value < 0.05. For the analysis of mouse lung samples, we excluded the neutrophil population.

#### Compositional analysis

To estimate the changes in the proportions of cell populations across experimental conditions we applied the *scCODA* package (version 0.1.2) (34), a Bayesian approach that considers inherent biases in cell-type compositions and addresses the low-replicate issue, making it an appropriate tool for our experimental design. Biological replicates were considered as covariates when available. This compositional analysis depends on a reference cell-type, and credible changes (nominal FDR < 0.05) for varying log-fold changes should be interpreted in relation to the selected reference cell-type. For this reason, we used the “automatic” selection of reference cell-type for single comparisons, but we set a manual “pre-selected reference” cell-type when multiple comparisons were conducted against the same condition. In the latter case, we tested multiple reference cells to ensure consistent results.

#### Differential expression analysis (DEA)

To define condition-specific signatures, we performed differential expression analysis (DEA) using the Seurat function “*FindMarkers*” with the MAST test, taking advantage of the latent variable correction in cases where biological replicates were present. We defined genes to be significantly differentially expressed (sDE) if the Log2 Fold Change (Log2FC) > |1|, with a False Discovery Rate (FDR) adjusted p-value < 0.05, and if they were present in at least 10% of cells.

#### Gene Set Enrichment analysis (GSEA)

To unravel biological pathways affected by a specific condition, we performed a pre-ranked GSEA with the *fgsea* package (35), and employed multiple gene sets from the Human Molecular Signatures Database (MSigDB); specifically, the mouse and human Hallmarks (H), Canonical Curated Pathways (C2:CP) such as Reactome, and Gene Ontology (C5:GO) gene sets from Biological Process (BP).

We excluded gene sets that did not satisfy our pathaway size criteria (10-300 genes) and considered significant only results with an FDR adjusted p-value < 0.05, where >5 genes overlapped with the gene set, and this accounted for >15% of the gene set.

#### Condition-specific signatures

To evaluate the effectiveness of the FixNCut protocol in preventing cells from undergoing stress, we assessed condition-specific signatures and sDE genes associated with external stressors. For human PBMC samples, we included publicly available signatures, such as the *ex-vivo* PBMC handling signature on microarrays by Baechler *et al.* (36) and the human PBMC sampling time-dependent signature on single-cell by Massoni-Badosa *et al.* (4). Additionally, we studied multiple dissociation-induced gene expression signatures for the tissue samples, including dissociation-associated on mouse muscle stem cells by van den Brink *et al.* (19), warm-dissociation on mouse kidney samples by Denisenko *et al*. (3), and warm collagenase-associated on mouse primary tumors and patient-derived mouse xenografts by O’Flanagan *et al*. (2). With this purpose, we downloaded the sDE genes from these studies and computed signature-specific scores using the *Ucell* package (37). To account for biases arising from differences in cell population numbers across protocols, we downsampled at a maximum of 250 cells per cell-type and replicate. Moreover, we also investigated the expression of specific genes across cell populations and conditions. Statistical analysis between sample protocols was performed using the Wilcoxon signed-rank test.

## Abbreviations

ACME: ACetic-MEthanol
CM: Complete Media
DEA: Differential Expression Analysis
DSP: Dithiobis(succinimidyl propionate)
DSP-PE: DSP-fixed, paraffin-embedded
FFPE: Formalin-fixed, paraffin-embedded
GEM: Gel Beads-in-emulsion
GSEA: Gene set enrichment analysis
H&E: Hematoxylin and Eosin
HBSS: Hank’s Balanced Salt Solution
HVG: Highly Variable Gene
IBD: Inflammatory Bowel Disease
LMO: Lipid-Modified Oligo
mAbs: monoclonal antibodies
ME: Mean Expression
MFI: Mean Fluorescent Intensity
NCP: Neural Cell Populations
NHS ester: N-hydroxysuccinimide ester
PBMC: Peripheral Blood Mononuclear cells
PC: Principal Components
QC: Quality Control
ROS: Reactive Oxygen Species
RT: Room Temperature
STEP: Spatial Tissue Exploration Program
UMAP: Uniform Manifold Approximation and Projection
UMI: Unique Molecular Identifier
UPR: Unfolded Protein Response

## Declarations

### Ethics approval and consent to participate

Experiments with mice were conducted in accordance with the guidelines of the Animal Care and Use Committee of Barcelona Science Park under protocol CEEA-PCB-14-000275. Healthy PBMC donors were collected with written informed consent at the Centro Nacional de Análisis Genómico (CNAG-CRG) with ethical approval granted by the local ethics committee. Healthy human buffy coats were provided by the Australian Red Cross Blood Service, and the ethical approval was obtained by the Melbourne Human Ethics Committee (approval 01/14). Colon biopsies from Ulcerative Colitis patients in remission were collected with written informed consent at the Hospital Clinic of Barcelona with ethical approval granted by the clinical research ethics committee of the Hospital Clínic of Barcelona. Prostate tissues were collected with written informed consent from previously-untreated patients undergoing radical prostatectomy at St. Andrew’s Hospital, Adelaide, Australia, through the Australian Prostate Cancer BioResource. Ethical approval for tissue collection and experimentation was obtained from St Andrew’s (approval #80) and the University of Adelaide (approval H-2012-016) Human Research Ethics committees. This study was conducted in accordance with the Declaration of Helsinki principles.

### Availability of data and materials

The complete raw data (FASTQ files) generated in this study have been submitted to the NCBI Gene Expression Omnibus (GEO). The count matrices and metadata have been deposited at Zenodo. All data will be available upon publication.

The code to reproduce the full analysis is hosted at Github: https://github.com/LJimenezGracia/FIXnCUT_benchmarking.

## Authors’ contributions

H.H. and L.G.M designed the study. L.G.M and J.P devised the use of DSP for fixing tissues following dissociation. D.M. and G.C. performed the scRNA-seq experiments. L.J.-G. and J.C.N. annotated all datasets. L.J.-G. performed the computational analysis. D.M., J.C.N., S.R., E.M-A. and V.G. performed and analyzed FACS data. K.W. and K.B.J. performed the CODEX experiments. H.-M. performed all mice work. N.K.R., L.M.B., J.P.B., F.T., L.K.S., S.S., M.v.d.B., T.K, P.L.v.d.V., M.C.N., P.R., A.S. provided clinical material. L.J.-G., D.M. and J.C.N. prepared the figures. L.J.-G, D.M, H.H. and L.G.M. wrote the manuscript with contributions from all the co-authors. All authors read and approved the current version of the manuscript.

## Acknowledgements

The LMO oligos used were kindly provided by Chris McGinnis from the Gartner lab (UCSF). The authors would like to thank the CRG Scientific IT Unit and the maintainers of the CNAG-CRG compute cluster for providing assistance with essential computing resources.

## Disclaimer

Content of this publication reflects only the author’s view and the JU is not responsible for any use that may be made of the information it contains.

