## Supplementary Material for "FixNCut: Single-cell genomics through reversible tissue fixation and dissociation"

### Supplementary Information

***Additional file 1: Supplementary material.***

*Supplementary Figures 1-6 and Supplementary figure and table legends.*

#### Additional Figures

**Additional file 1: Figure S1. FixNCut protocol tested in human PBMCs.** (**a**) Bright-field microscopy images of fresh (**top**) and fixed (**bottom**) human PBMCs. (**b**) cDNA profiles of human PBMCs in fresh (**top**) and fixed (**bottom**) samples. (**c**) Final GEX library BioA profiles of human PBMCs in fresh (**top**) and fixed (**bottom**) samples. (**d**) Comparative analysis of the number of detected UMIs (**left**) and genes (**right**) for each cell barcode based on sequencing reads using a linear model. (**e**) Distribution of main QC metrics (number of UMIs, genes, and percentage of mitochondrial genes) by protocol. (**f**) Distribution of main QC metrics by cell type and protocol. (**g**) Overlap of highly variable genes (HVGs) shared between fresh and fixed human PBMCs, considering 3000 HVGs computed for each sample independently. (**h**) Linear regression model comparing the average gene expression levels of expressed genes by cell type. The coefficient of determination (R2) computed with Pearson correlation and the corresponding p-value are indicated. (**i**) Dot plot showing the average expression for top genes (*x-axis*) across all 19 cell types (*y-axis*). Dot size represents the percentage of cells in a cluster expressing each gene, and the color indicates the average expression level. (**j**) Score of sampling-time gene signature (4) for fresh and fixed human PBMCs across cell populations. Statistical analysis between fixed and fresh cells was performed using Wilcoxon signed-rank test; significance results are indicated (ns, p>0.05, * p<=0.05, ** p<=0.01, *** p<=0.001, **** p<=0.0001). (**k**) Overlap between significant DE genes enriched in fresh PBMCs with genes from the sampling-time signature (4).

**Additional file 1: Figure S2. FixNCut protocol tested in mouse lung samples.** (**a**) Bright-field (**left**), DAPI fluorescent (**middle**), and two microscopy images overlay (**right**) of fresh (**top**) and fixed (**bottom**) mouse lung cells. (**b**) UpSet plot showing the intersection between the total captured genes in each library; highlighting genes detected in all libraries (blue), and exclusive to fixed (orange) and fresh (green) samples. (**c**) Comparative analysis of the number of detected UMIs (**left**) and genes (**right**) for each cell barcode based on sequencing reads using a linear model. (**d**) Distribution of the main QC metrics (number of UMIs, genes, and percentage of mitochondrial genes) by protocol, considering all cell types (**top**) or after neutrophil exclusion (**bottom**). (**e**) Distribution of the main QC metrics by cell-type and protocol. (**f**) Overlap of highly variable genes (HVGs) shared between fresh and fixed mouse lung samples, considering 3000 HVGs computed for each replicate independently (**bottom**) or by protocol (**top**). (**g**) UMAP visualization comparison across protocol and replicates. (**h**) Linear regression model comparing average gene expression levels of expressed genes between fresh and fixed samples by cell-type. The coefficient of determination (R2) computed with Pearson correlation and the corresponding p-value are indicated. (**i**) Dotplot showing average expression for the top genes (*x-axis*) for all 20 cell types (*y-axis*). The dot size reflects the percentage of cells in a cluster expressing each gene, and the color represents the average expression level.

**Additional file 1: Figure S3. FixNCut protocol tested in mouse colon samples.** (**a**) Comparative analysis of the number of detected UMIs (**left**) and genes (**right**) for each cell barcode based on sequencing reads using a linear model. (**b**) Distribution of main QC metrics (number of UMIs, genes, and percentage of mitochondrial genes) by protocol. (**c**) Distribution of main QC metrics by cell type and protocol. (**d**) Overlap of highly variable genes (HVGs) shared between fresh and fixed mouse colon samples, considering 3000

HVGs computed for each sample independently. (**e**) Linear regression model comparing the average gene expression levels of expressed genes by cell type. The coefficient of determination (R2) computed with Pearson correlation and the corresponding p-value are indicated. (**i**) Dotplot showing average expression for the top genes (*y-axis*) for all 16 cell types (*x-axis*). The dot size reflects the percentage of cells in a cluster expressing each gene, and the color represents the average expression level.

**Additional file 1: Figure S4. Long-term storage of fixed mouse lung samples.** (**a**) Comparative analysis of the number of detected UMIs (**left**) and genes (**right**) for each cell barcode based on sequencing reads using a linear model. (**b**) Distribution of main QC metrics (number of UMIs, genes and percentage of mitochondrial genes) by protocol, considering all cell types (**top**) or after neutrophil exclusion (**bottom**). (**c**) Distribution of main QC metrics by cell-type and protocol. (**d**) Overlap of highly variable genes (HVGs) shared between protocols, considering the 3000 HVGs computed for each protocol independently. (**e**) Linear regression model comparing the average gene expression levels of expressed genes by cell-type and protocol; fixed+cryo vs fixed (**top**) and fixed+cryo vs cryo (**bottom**). The coefficient of determination (R2) computed with Pearson correlation and the corresponding p-value are indicated. (**f**) Dotplot showing average expression for the top genes (x-axis) for all 20 cell types (y-axis). Dot size reflects the percentage of cells in a cluster expressing each gene, and the color represents the average expression level.

**Additional file 1: Figure S5. FixNCut protocol tested in human colon biopsies.** (**a**) Comparative analysis of the number of detected UMIs (**left**) and genes (**right**) for each cell barcode based on sequencing reads using a linear model. (**b**) Distribution of main QC metrics (number of UMIs, genes and percentage of mitochondrial genes) by protocol. (**c**) Distribution of main QC metrics by cell-type and protocol. (**d**) Overlap of highly variable genes (HVGs) shared between protocols, considering the 3000 HVGs computed for each condition independently. (**e**) Linear regression model comparing average gene expression levels of expressed genes by cell-type and protocol; fixed vs fresh (**top**) and fixed+cryo vs cryo (**bottom**). (**f**) Dotplot showing average expression for the top genes (*x-axis*) for all 21 cell types (*y-axis*). Dot size reflects the percentage of cells in a cluster expressing each gene, and the color represents the average expression level. (**g**) Score of warm-dissociation gene signature (3) for human mouse colon by cell population across protocols.

**Figure S6. Flow cytometry analysis in fixed cells and tissues in mouse and human.** (**a**) Representative flow cytometry experiment of forward scatter (FSC) and side scatter (SSC) in fresh (**left**) and fixed (**right**) cryopreserved human PBMCs from healthy donors (n=3). (**b**) Representative flow cytometry experiment of cell viability based on DAPI expression in fresh (**left**) and fixed (**right**) cryopreserved human PBMCs from healthy donors (n=3). (**c**) Representative flow cytometry experiment of cell apoptosis based on the expression of Annexin V FITC and PI in fresh (**left**) and fixed freshly isolated PBMCs (**right**). Cells were analyzed after isolation and fixation at day 0 (**top**) and 2-days after fixation (**bottom**) (n=4). (**d**) Representative gating strategy of one experiment analyzed by flow cytometry of freshly isolated PBMCs from healthy donors (n=4). PBMCs were stained with anti-human CD3, CD19, CD4 and CD8 monoclonal antibodies (mAbs). T cells were selected by the positive expression of CD3. CD4 positive and CD8 positive T cells were selected from CD3 positive. (**e**) Representative flow cytometry experiment of forward scatter (FSC) and side scatter (SSC) (**top**) and cell viability based on DAPI expression (**bottom**) of human colon biopsies from IBD donors.

#### Additional Tables

**Additional file 2: Table 1.**

This table contains the differentially expressed (DE) genes between fixed and fresh human peripheral blood mononuclear cells (PBMCs), considering all cell populations. The differential expression analysis was performed using Seurat MAST, and genes with an FDR adjusted p-value < 0.05 and present in at least 10% of cells were considered DE (see Methods). Genes with positive Log2FC indicate higher expression in fixed PBMCs, while those with negative Log2FC are DE in fresh PBMCs.

**Additional file 3: Table 2.**

This table contains differentially expressed (DE) genes between fixed and fresh human PBMCs, separated by cell population. The analysis was performed using Seurat MAST, with genes considered as DE with a FDR adjusted p-value < 0.05 and present in at least 10% of cells (see Methods). A positive Log2FC value indicates higher expression in fixed PBMCs, while a negative value indicates differential expression in fresh PBMCs.

**Additional file 4: Table 3.**

The table contains the distribution statistics of the mean gene expression for detected genes of mouse lung samples, including those detected in any of the four libraries (2 fixed and 2 fresh replicates), those detected in all libraries (considering cells from both protocols together and independently), and genes exclusively detected in both replicates from fixed or fresh conditions. These statistics provide insights into the quality and consistency of gene capture across protocols and replicates.

**Additional file 5: Table 4.**

This table contains differentially expressed (DE) genes for mouse tissue comparisons, considering all cell populations. The comparisons include: 1) fixed versus fresh lung samples, 2) fixed versus fresh colon samples, 3) fixed compared to cryopreserved (cryo), 4) fixed+cryo versus cryo, and 5) fixed+cryo compared with fixed on lung samples. The statistical analysis was performed using Seurat MAST with genes considered DE if they had an FDR adjusted p-value < 0.05 and were present in at least 10% of cells (see Methods). The results include positive Log2FC values indicating higher expression in the first condition mentioned, while negative values indicate higher expression in the second condition mentioned.

**Additional file 6: Table 5.**

This table contains the list of differentially expressed (DE) genes between fixed and fresh mouse lung samples, categorized by cell population. The test was performed using Seurat MAST, where genes with a FDR adjusted p-value < 0.05 and present in at least 10% of cells were considered as DE (see Methods). The Log2FC values indicate the direction of expression change, where positive values indicate higher expression in fixed samples and negative values indicate higher expression in fresh samples.

**Additional file 7: Table 6.**

This table contains the results of gene set enrichment analysis (GSEA) between fixed and fresh mouse lung samples by cell population. The analysis was performed using the *fgsea* package and multiple gene sets from MSigDB. Gene sets containing 10-300 genes were included, and only results with a FDR adjusted p-value < 0.05 were considered if they contained >5 overlapping genes, which represents >15% of the gene set (see Methods).

**Additional file 8: Table 7.**

This table contains the differentially expressed (DE) genes between fixed and fresh mouse colon samples, analyzed by cell population using Seurat MAST. Genes were considered DE if they had an FDR adjusted p-value < 0.05 and were present in at least 10% of cells (see Methods). The Log2FC value indicates whether the gene is more highly expressed in fixed (positive values) or fresh (negative values) samples.

**Additional file 9: Table 8.**

This table contains the results of gene set enrichment analysis (GSEA) between fixed and fresh mouse colon samples by cell population. The analysis was performed using the *fgsea* package and multiple gene sets from MSigDB. Gene sets containing 10-300 genes were included, and results were considered significant if they had an FDR adjusted p-value < 0.05 and if >5 genes from the gene set overlapped with the differentially expressed genes, representing more than 15% of the gene set (see Methods).

**Additional file 10: Table 9.**

This table contains differentially expressed (DE) genes between fixed+cryo and cryo mouse lung samples, analyzed by cell population. The test was performed with Seurat MAST, and genes were considered DE with a FDR adjusted p-value < 0.05, and present in at least 10% of cells (see Methods). A positive Log2FC indicates higher expression in fixed or fixed+cryo samples, whereas negative values indicate differential expression in cryo or fixed samples.

**Additional file 11: Table 10.**

This table contains the results of gene set enrichment analysis (GSEA) for fixed+cryo versus cryo mouse lung samples, by cell population. The analysis was performed using the *fgsea* package with multiple gene sets from MSigDB. Gene sets containing within 10-300 genes were included and the obtained results were considered significant if the FDR adjusted p-value was <0.05 and if >5 genes overlapped with the gene set, representing >15% of the gene set (see Methods).

**Additional file 12: Table 11.**

This table contains differentially expressed (DE) genes for human colon biopsies comparisons, considering all cell populations. The comparisons include: 1) fixed and fresh, 2) fixed and cryo, 3) fixed+cryo versus cryo, and 4) fixed+cryo compared with fixed samples. The analysis was performed using Seurat MAST, and genes were considered DE with a FDR adjusted p-value < 0.05 and present in at least 10% of cells (see Methods). A positive Log2FC indicates higher expression in fixed or fixed+cryo samples, whereas negative values indicate differential expression in fresh, cryo or fixed samples.

**Additional file 13: Table 12.**

This table contains the differentially expressed (DE) genes between fixed and fresh human colon biopsies, separated by cell population. The test was performed using Seurat MAST, with genes considered DE if they had a FDR adjusted p-value < 0.05 and were present in at least 10% of cells (see Methods). Positive Log2FC values indicate higher expression in fixed samples, while negative values indicate higher expression in fresh samples.

**Additional file 14: Table 13.**

This table contains the differentially expressed (DE) genes between fixed+cryo versus cryo human colon biopsies, separated by cell population. The test was performed using Seurat MAST, with genes considered DE if they had a FDR adjusted p-value < 0.05 and were present in at least 10% of cells (see Methods). Positive Log2FC values indicate higher expression in fixed+cryo samples, while negative values indicate higher expression in cryo samples.

**Additional file 15: Table 14.**

This table contains the results of gene set enrichment analysis (GSEA) for fixed versus fresh human colon biopsies comparison by cell population. The test was performed with the *fgsea* package using multiple gene sets from MSigDB. Gene sets were included only if they contained between 10-300 genes, and obtained results were considered with a FDR adjusted p-value < 0.05, only if more than 5 genes were overlapping with the gene set and represented more than 15% of the gene set (see Methods).

**Additional file 16: Table 15.**

This table contains the results of gene set enrichment analysis (GSEA) for fixed+cryo versus cryo human colon biopsies comparison by cell population. The test was performed with the *fgsea* package using multiple gene sets from MSigDB. Gene sets were included only if they contained between 10-300 genes, and obtained results were considered with a FDR adjusted p-value < 0.05, only if more than 5 genes were overlapping with the gene set and represented more than 15% of the gene set (see Methods).

Figure S1

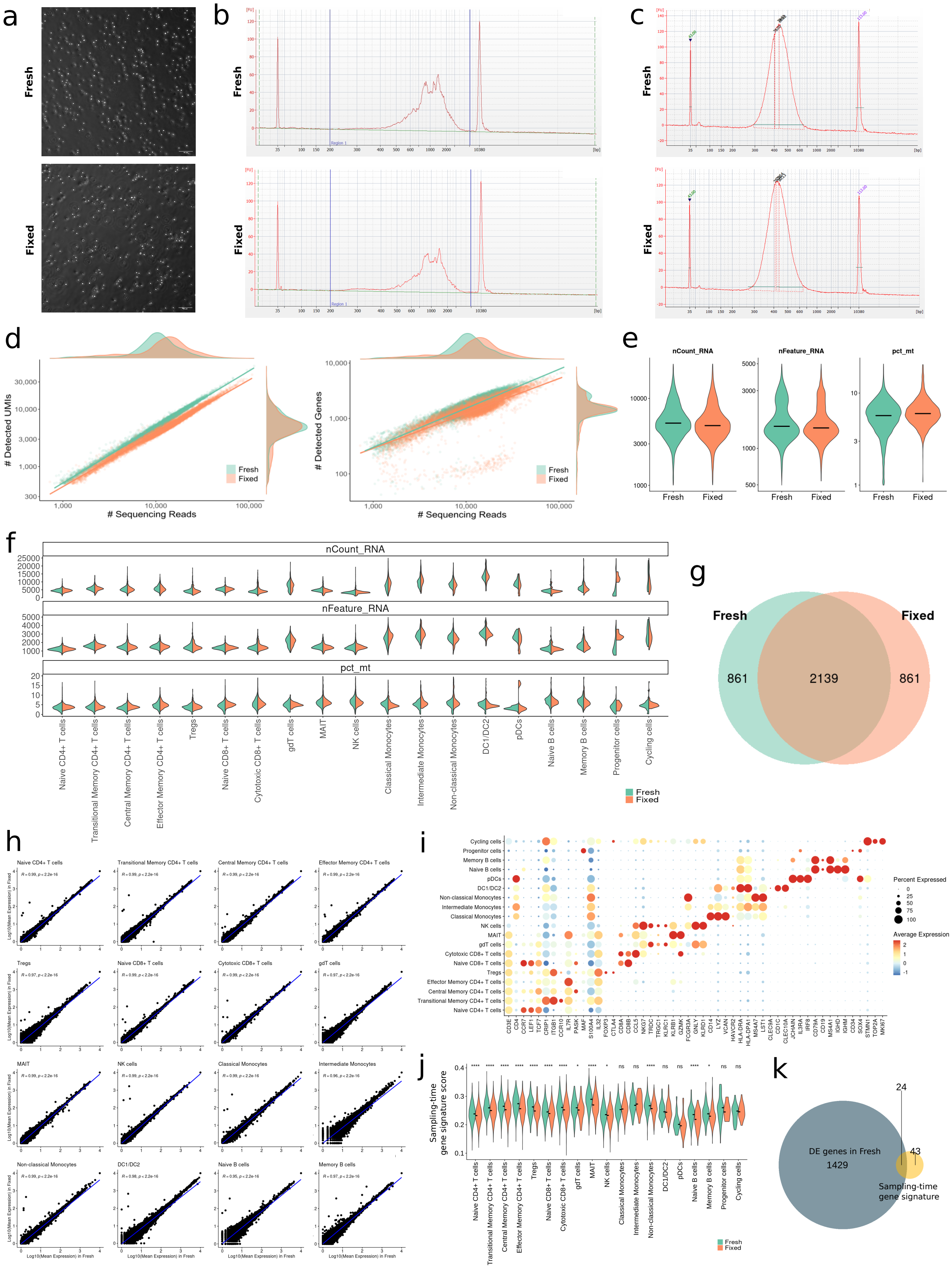

a
Fresh
Fixed
b
Fresh
Fixed
c
Fresh
Fixed
d
### Detected UMIs
### Sequencing Reads
### Detected Genes
Fresh
Fixed
e
nCount_RNA
nFeature_RNA
pct_mt
Fresh
Fixed
f
nCount_RNA
nFeature_RNA
pct_mt
Naive CD4+ T cells
Transitional Memory CD4+ T cells
Central Memory CD4+ T cells
Effector Memory CD4+ T cells
Tregs
Naive CD8+ T cells
Cytotoxic CD8+ T cells
gdT cells
MAIT
NK cells
Classical Monocytes
Intermediate Monocytes
Non-classical Monocytes
DC1/DC2
pDCs
Naive B cells
Memory B cells
Progenitor cells
Cycling cells
Fresh
Fixed
g
Fresh
Fixed
861
2139
861
h
Log10(Mean Expression) in Fixed
Log10(Mean Expression) in Fresh
i
Percent Expressed
Average Expression
j
Sampling-time gene signature score
Fresh
Fixed
k
DE genes in Fresh
1429
24
43
Sampling-time gene signature

Figure S2

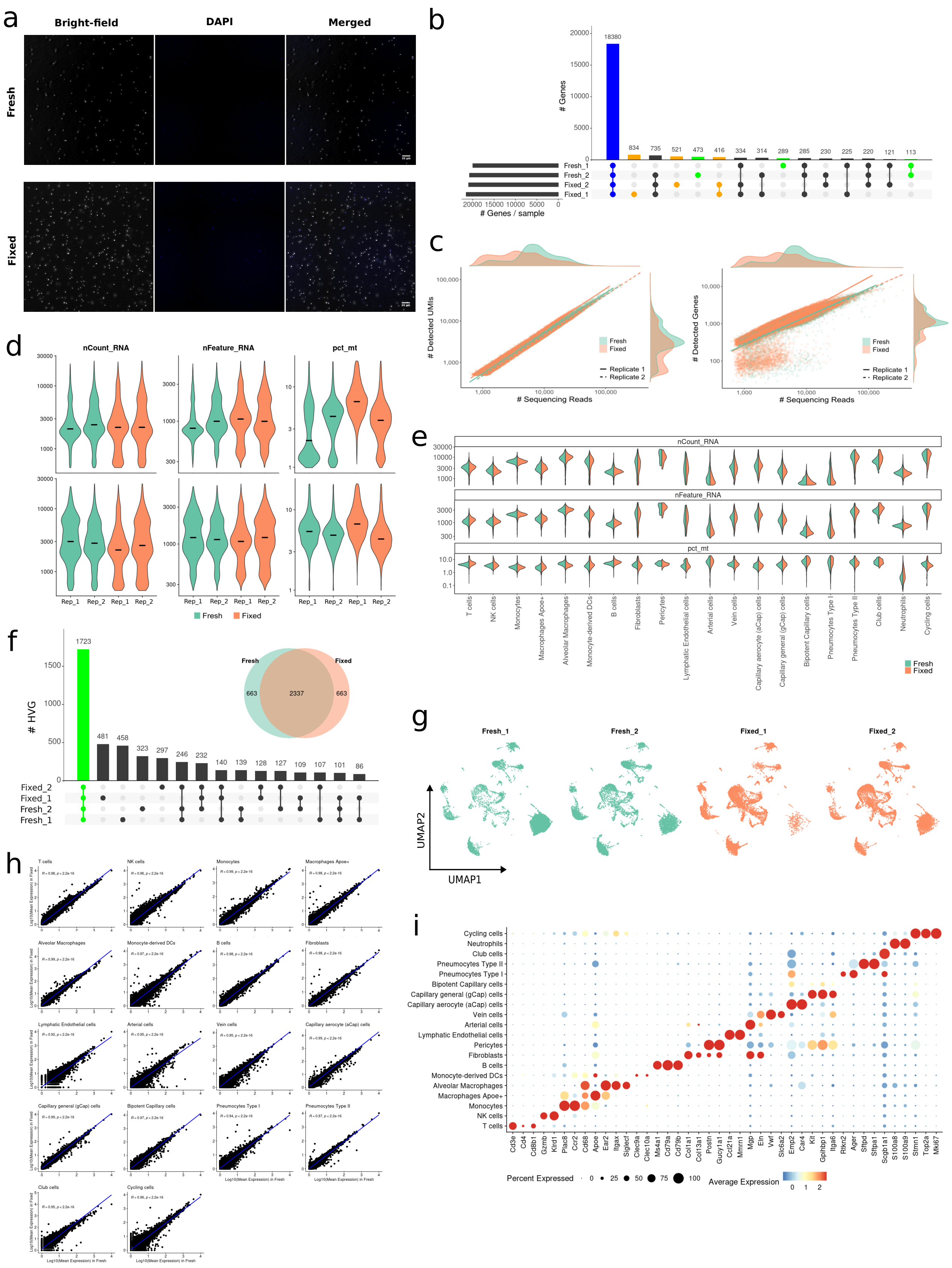

Figure S3

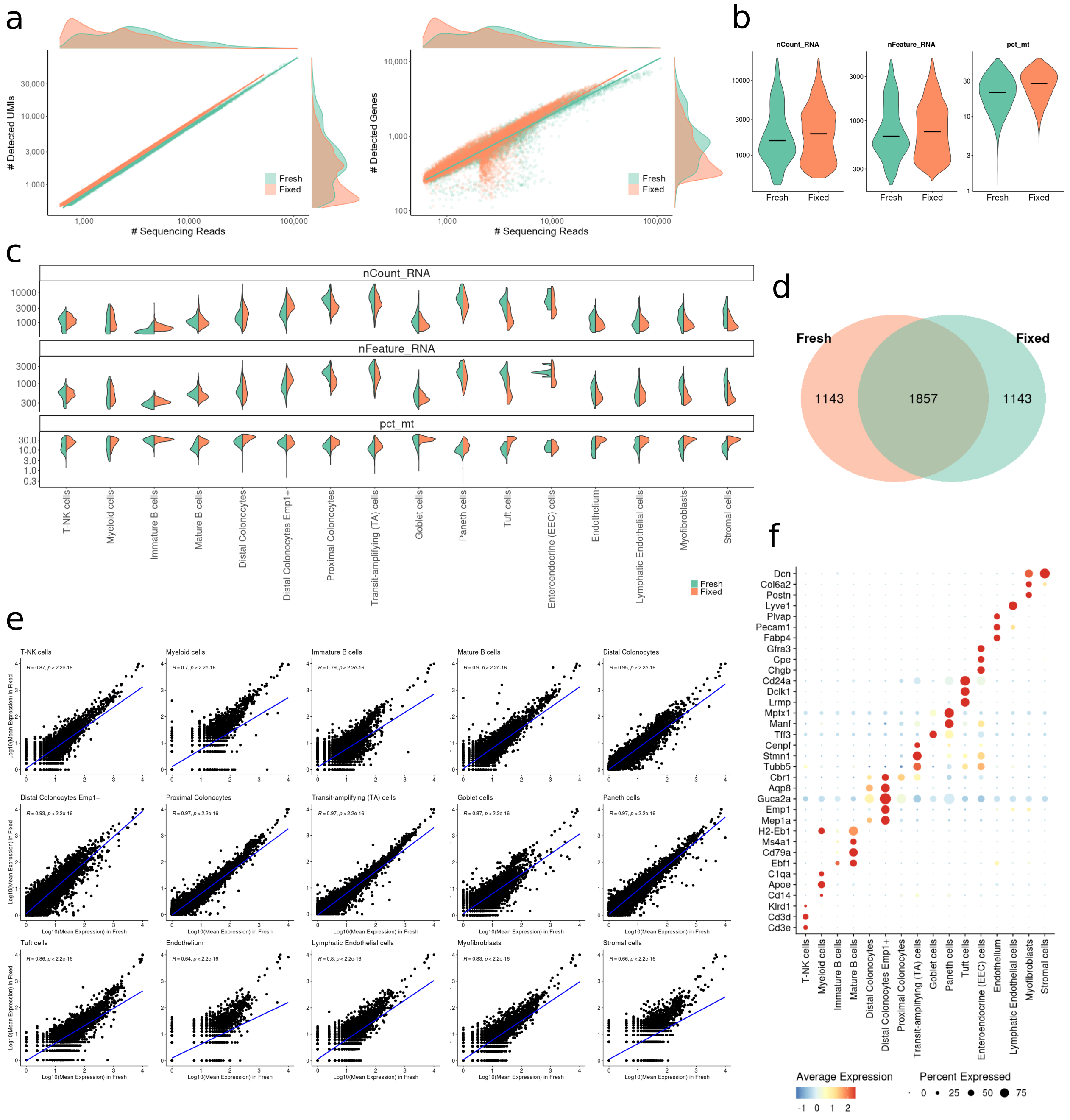
a
### Detected UMIs
### Sequencing Reads
### Detected Genes
Fresh
Fixed
b
nCount_RNA
nFeature_RNA
pct_mt
c
d
Fresh
Fixed
1143
1857
1143
e
f
T-NK cells
Myeloid cells
Immature B cells
Mature B cells
Distal Colonocytes
Distal Colonocytes Emp1+
Proximal Colonocytes
Transit-amplifying (TA) cells
Goblet cells
Paneth cells
Tuft cells
Enteroendocrine (EEC) cells
Endothelium
Lymphatic Endothelial cells
Myofibroblasts
Stromal cells
Log10(Mean Expression) in Fresh
Log10(Mean Expression) in Fixed
Average Expression
Percent Expressed

Figure S4

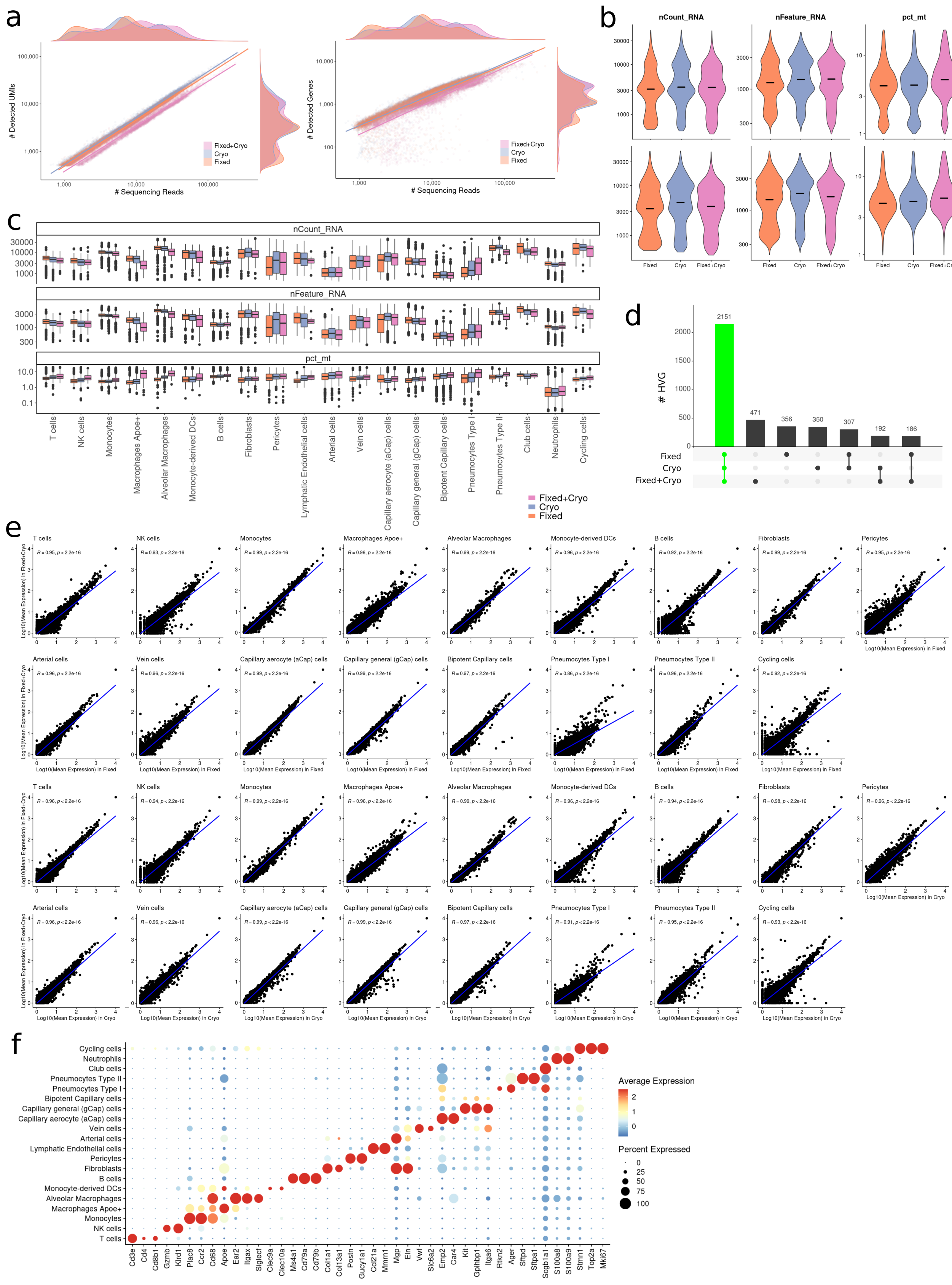
a
### Detected UMIs
### Sequencing Reads
Fixed+Cryo
Cryo
Fixed
### Detected Genes
b
nCount_RNA
nFeature_RNA
pct_mt
c
d
### HVG
Fixed
Cryo
Fixed+Cryo
e
Log10(Mean Expression) in Fixed+Cryo
Log10(Mean Expression) in Fixed
Log10(Mean Expression) in Cryo
f
Average Expression
Percent Expressed
T cells
NK cells
Monocytes
Macrophages Apoe+
Alveolar Macrophages
Monocyte-derived DCs
B cells
Fibroblasts
Pericytes
Lymphatic Endothelial cells
Arterial cells
Vein cells
Capillary aerocyte (aCap) cells
Capillary general (gCap) cells
Bipotent Capillary cells
Pneumocytes Type I
Pneumocytes Type II
Club cells
Neutrophils
Cycling cells

Figure S5

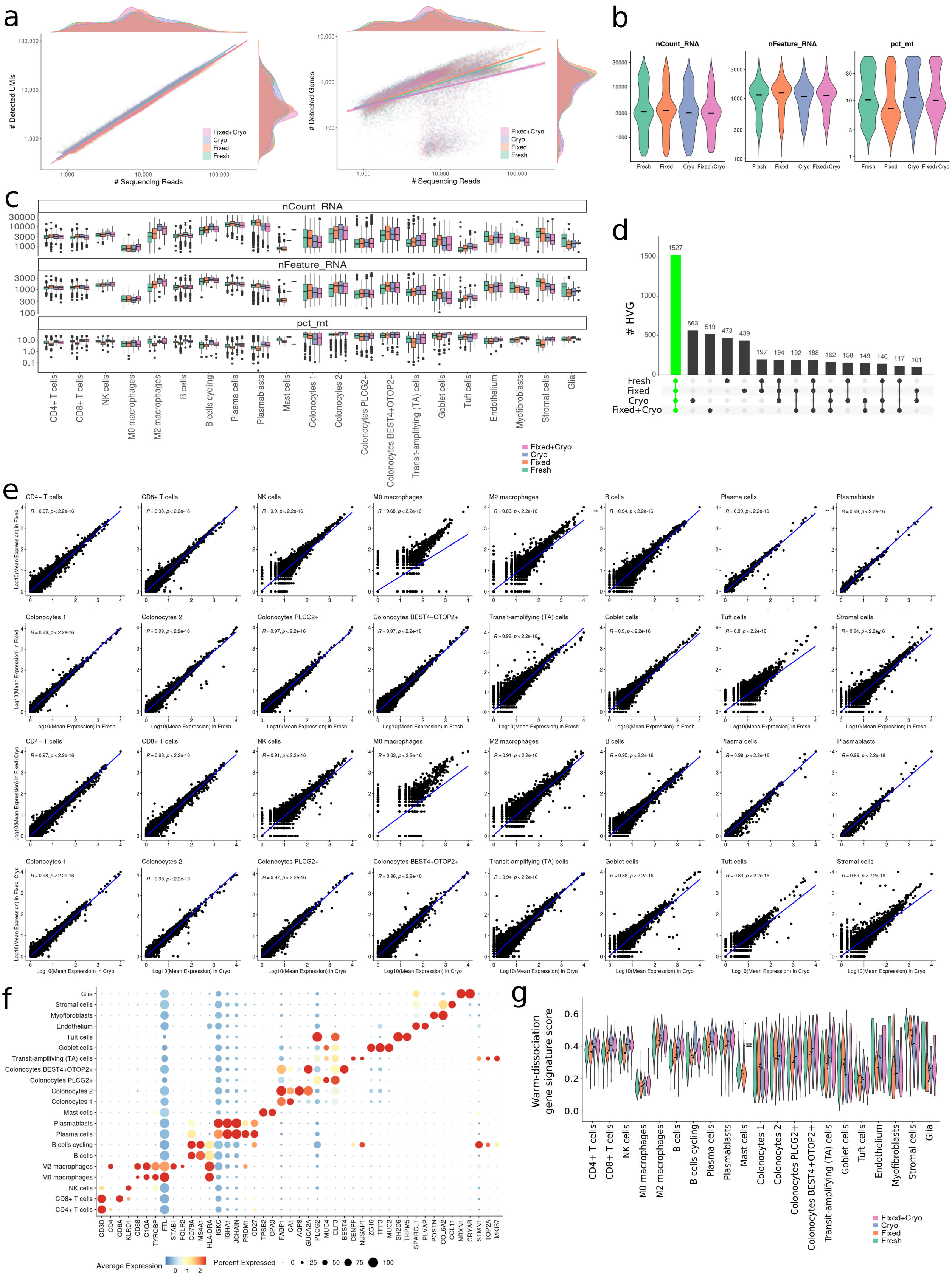
a
### Detected UMIs
### Sequencing Reads
### Detected Genes
Fixed+Cryo
Cryo
Fixed
Fresh
b
nCount_RNA
nFeature_RNA
pct_mt
Fresh
Fixed
Cryo
Fixed+Cryo
c
nCount_RNA
nFeature_RNA
pct_mt
CD4+ T cells
CD8+ T cells
NK cells
M0 macrophages
M2 macrophages
B cells
B cells cycling
Plasma cells
Plasmablasts
Mast cells
Colonocytes 1
Colonocytes 2
Colonocytes PLCG2+
Colonocytes BEST4+OTOP2+
Transit-amplifying (TA) cells
Goblet cells
Tuft cells
Endothelium
Myofibroblasts
Stromal cells
Glia
d
### HVG
1527
563
519
473
439
197
194
192
188
162
158
149
146
117
101
e
Log10(Mean Expression) in Fixed
Log10(Mean Expression) in Fresh
Log10(Mean Expression) in Fixed+Cryo
Log10(Mean Expression) in Cryo
f
Average Expression
Percent Expressed
g
Warm-dissociation gene signature score

Figure S6

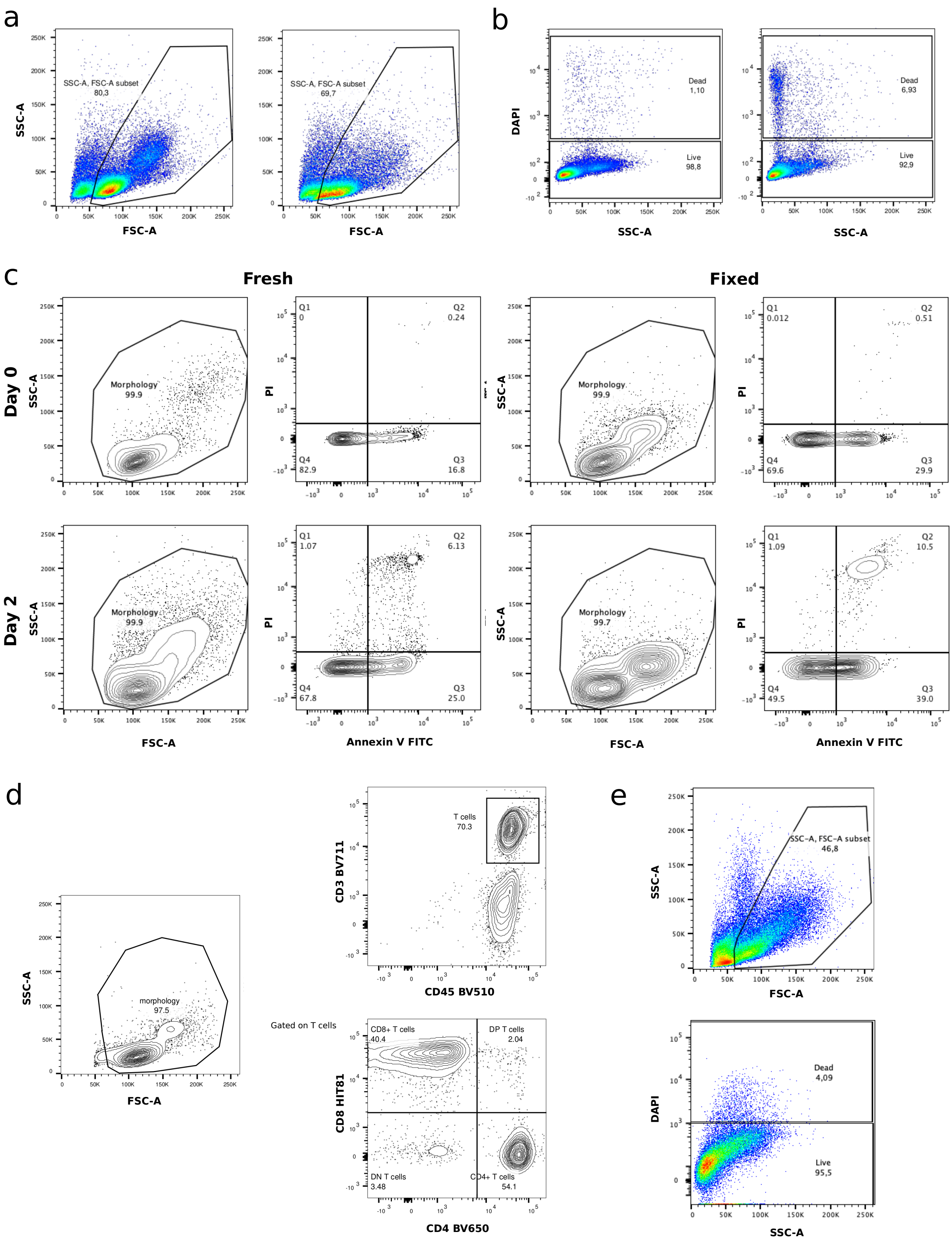
a
SSC-A, FSC-A subset
80,3
SSC-A, FSC-A subset
69,7
SSC-A
FSC-A
FSC-A
b
DAPI
Dead
1,10
Live
98,8
Dead
6,93
Live
92,9
SSC-A
SSC-A
c
Fresh
Fixed
Day 0
Day 2
SSC-A
Morphology
99.9
PI
Q1
0
Q2
0.24
Q4
82.9
Q3
16.8
Morphology
99.9
Q1
0.012
Q2
0.51
Q4
69.6
Q3
29.9
Morphology
99.9
Q1
1.07
Q2
6.13
Q4
67.8
Q3
25.0
Morphology
99.7
Q1
1.09
Q2
10.5
Q4
49.5
Q3
39.0
FSC-A
Annexin V FITC
FSC-A
Annexin V FITC
d
SSC-A
morphology
97.5
FSC-A
CD3 BV711
T cells
70.3
CD45 BV510
Gated on T cells
CD8 HIT81
CD8+ T cells
40.4
DP T cells
2.04
DN T cells
3.48
CD4+ T cells
54.1
CD4 BV650
e
SSC-A
SSC-A, FSC-A subset
46,8
FSC-A
DAPI
Dead
4,09
Live
95,5
SSC-A
